# A Hierarchical Model of the Hypothalamo-Adreno-Pituitary system

**DOI:** 10.1101/837724

**Authors:** Camilo La Rota, Dan Tudor Vuza, Adriana Climescu-Haulica

## Abstract

We present a model of the Hypothalamo-Adreno-Pituitary (HPA) system; our approach is hierarchical and biologically sound, reflecting the complexity of the real HPA system. The structure of the conceptual model is based on theoretical frameworks of biological complex systems and its implementation takes advantage of recent hybrid automata modelling and analysis tools, being modular and reflecting parameter and mechanisms uncertainties.

## 1 Introduction

The HPA system is a neuro-endocrine system whose activity is fundamental for the good functioning of multiple physiological functions. Shortly, the system is composed of: i) neuro-endocrine cell populations in the Hypothalamic Paraventricular Nucleus (PVN) whose non-myelinated axons have terminals in the median eminence (ME), which release Corticotropin Releasing Hormone (CRH), vasopressin (AVP) and other hormones into the hypophyseal portal venous system; ii) endocrine population cells (corticotrophes) at the anterior pituitary, which secrete adrenocorticotropic hormone (ACTH); iii) cortisol producing population cells of the Adrenal Cortex (ACx); and iv) other neural and pituitary cell populations.

The HPA shows multiple behaviors, among which the most evidents are circadian and ultradian oscillations or pulses. Several models have been proposed in the litterature (refs), each one focusing on one aspect of the system behavior, which have shed some light on some of the mechanisms underlying them. However, several important questions remain unanswered concerning the HPA behavior and its underlying dynamics: i) which is the origin of the ultradian pulses or oscillations, are they locally generated by the internal dynamics of a specific cell population, independently of the interactions among cell populations ? do they come from an external source ? or do they result from the endogenous dynamics of the HPA cell population network ? (and indeed, are they pulses or oscillations ?) ii) which is the mechanism and the influence of the circadian rhythms and light in the HPA behavior ? iii) which is or are the mechanisms leading to bistability (or multistability?) and which is its physiological meaning. iv) How do physical and psychological stress, as well as immunitary challenge, influence HPA dynamics ? v) How do other systems interact with the HPA system ?

*Neuroendocrine systems models:* A sequence of HPA subsystem models was developed by Keenan et al in [87, 88, 86], based on a previous model of the HPG (gonadal) subsystem presented in [90, 89]. The initial model involves the synthesis, acummulation and release mechanisms of hypothalamic CRH-AVP neurons, the ACTH corticotrops in the AP and the cortisol secreting cells of the ACx. A more detailed model of the AP (ACTH) → ACx (cortisol) interaction, involving kinetics of exchange of plasma cortisol among free and protein bound compartements was decribed in [88], and a still more detailed model of ACTH secretion based on these previous models was presented in [86]. An important feature of these models is the use of a stochastic pulsatory mechanism for the generation of CRH-AVP secretory pulses, which implicitly assumes that pulse generation is independent of the network dynamics; in addition, no difference is made between CRH and AVP dynamics and between their effects on corticotroph activity. This model is capable of a very faithfull reconstruction of hormone time-series and allows estimation and detailed statistical analysis. However, it has many free parameters, it is not easily amenable to mathematical analysis, nothing is known about its general dynamics and behavior and some features and parameters (such as the pulse generation mechanism and parameters) lack a clear biological interpretation. Alternatively, mathematical models based on simplified descriptions of endocrine cell functions [102, 95, 82, 33, 67], led to detailed mathematical analyses that allow the study of the general behavior of the system and provide mechanistic explanations of well known behaviors. Some of these modeling analyses suggest that ultradian oscillations are endogenously produced by the HPA system dynamics[102, 95, 82] and provide alternative hypothetical mechanisms; some others suggest focus on homeostasis and long term dynamical regime changes, suggesting putative mechanisms underlying both healthy and abnormal basal behavior [33, 67]. However, none of these models is capable of retrieving the different behaviors observed at different time scales; each one of these models lack some features that may be important in determining the dynamics of the system, such as intracellular secretion products accumulation before release, pulse-stimulated secretion and others. Finally, a more physiologically plausible model of corticotroph electrophysiological activity has been developped by LeBeau et al. [97, 98, 150] with the aim to integrate it into a complete model of corticotroph secretion. This model is biologically plausible, mathematically tractable, and has relatively few free parameters; however, is limited to phenomena ocurring at the very fine time and space scale of the cell membrane. Therefore, there’s still the need of a more comprehensive model of the HPA system which includes clearly identifiable and biologically pertinent parameters, at different scales, and at the same time with a simple mathematical description that is amenable to analysis and simulation.

*Complex Biological Modelling approaches:* given the underlying complexity of biological systems, the construction and integration of realistic models need to be based on sound theoretical bases in order to reduce the complexity of the implementation and to facilitate analysis. We want to mention two theoretical frameworks that go in that direction. First, the mathematical theory of integrative physiology (*MTIP*) developed by G. Chauvet, in which he proposes to describe a complex biological system by decomposing it on several smaller models all of them based on the same kind of formalization such that they can be studied with the same mathematical tools and can then be reassembled to study the integrated system by mathematical analysis and computational simulations. The composing models live at different temporal and spatial scales, but retain the same mathematical structure. Models of neural systems based on Chauvet’s Mathematical Theory of Integrative Physiology [24] involving neurotransmitter release and synaptic phenomena have been proposed by [12, 13, 25]. While these articles are not about hormonal systems, they are interesting from the methodological point of view because they illustrate the application of the MTIP modeling framework, which is proposed by Chauvet as a standard way to construct complex models of physiological systems. Two different computer implementations of this modeling methodology have been shown in [13] and [99, 124]. In addition, neural and hormonal systems share many features, such as the secretion of chemical substances by endocrine and neuroendocrine cell populations in order to send information to other cell populations, as well as the membrane excitability and the generation of action potentials. Second, we must mention Mesarovic et al proposal [118] of using the complex systems modelling framework (systems of systems) in order to study how the categories of systems are organized in order to achieve a function under changing environmental conditions. The aim of both approaches is the search for organizing principles for biology, and as a result they give us ideas on how a system may be organized and therefore on how a computer model may also be organized in order to reflect the activity and function of the real biological system.

*Hybrid automata:* some important implementation problems found when modelling biological systems are: i) the lack of knowledge about the precise mechanisms underlying dynamical behavior as well as about precise values for many parameters of the system; ii) the ubiquitous prescence of nonlinearities and the complexity of the systems under study that make the analysis of their dynamics by mathematical methods very difficult; iii) this same complexity hampers our ability to study the system by computational methods, because of the difficulty to track numerical errors, the computational burden that implies the numerical analysis of the parameter space and of the behavioral space, and because of the difficult problem of software design. This same problems arise in other fields, such as in the analysis of embedded systems, and new methods and tools have been developped in order to help to solve them. Hybrid automata is a relatively new modelling paradigm that was introduced in order to describe, simulate and study real world systems consisting of both real valued state variables continuosly changing and discrete control variables abruptly changing the dynamics of the system [7, 73, 111]; this paradigm seems to be suitable for the modelling of biological systems due to the existence of strong nonlinearities in biological mechanisms that change abruptly the dynamics of subsystems in response to small variations of their inputs, or as a result of changes in their parameters due to changes in the behavior of other related systems. Model checking, reachability analysis, time verification, and parameter scanning methods and tools for the analysis of complex systems, and in particular for hybrid automata [74, 58, 59], have also been recently developped and applied succesfully to real-world nonlinear and complex systems [57, 43]. Applications in systems biology begin to emerge [133, 55, 11, 119, 32, 39].

*Our model:* We focus on a detailed corticotroph model embedded in a simplified HPA network, trying to be faithfull to current knowledge of corticotroph biology and electrophysiology (a bottom up approach); as our aim is to construct an integrated model of the HPA system, we make important simplifications to take into account only dynamical behavior ocurring from the minutes up to hours time-scales. Our approach is modular and hierarchical, the complete model is composed of various categories of subsystems, each subsystem corresponding to a specific functional unit acting at a given time-scale and living in a given structural unit at a given space-scale. This complex system structure is implemented as a hierarchical set of hybrid automata and implemented in the SpaceEx tool [59], which allows the composition of large, hierarchically organized sets of linear hybrid automata. Reachability analysis is currently being performed for the fast secretion control subsystem of the corticotroph. The next step is to include other modules and close the simplified HPA circuit in order to study the closed circuit dynamics.

## 2 Methods

### 2.1 Modelling framework

Our approach borrow ideas from the MTIP theoretical framework developed by Chauvet [23, 24] and from complex systems theory approaches to biological systems and hierarchical organization [118]. Some common fundamental concepts of both theoretical frameworks are those of *multilevelness* and *hierarchy*, and the existence of *organizing principles* increasing the stability domain of the global system [118, 23]. Biological systems have a double hierarchical organization, the functional and structural hierarchy [23, 24]. *Functional interactions* are non-local and non-symmetric, they represent the action or influence of an entity onto another by means of a *signal* flowing from the first one (source) to the second one (sink, or target); signals are transformed at the target, which is itself a source for other functional interactions. A physiological function is composed of functional interactions, acting at a given time scale, and different physiological functions of the same system interact across the temporal hierarchy, each one retaining its individuality and own behavior [23, 24]. A concept that is central in physics and dynamic systems theory but does not appear explicitly in the MITP is the *state* of an entity or of a system, the concept of *functional interaction* being used both to represent the *signal* emited from an entity to another and representing the *state* [24], due to the continuous hierchical spaces representation (there are no individual units but units continuous distributions. In our approach these two concepts, *state* and *signal* describe different observables.

**Rationale:**

1. The HPA system is a complex system and it is therefore composed of interconnected and interacting subsystems, each subsystem having its own behavior and being itself composed either of interacting subsubsystems or elements. Each subsystem is be represented by a directed graph (*G*_*l*_), where each vertex is an entity (or subsystem) and each arc a functional interaction.
2. Each entity (system, subsystem, subsubsytem, …, elementary entity) is characterized by a variable, the *state* (*χ*_*i*_), which represents the instantaneous condition of the entity *i* (e.g. excitability of a cell), it changes continuously depending on dynamic forces either applied to the entity or being internally generated; it is a property of the entity and can be a scalar or a vector variable.
3. Each entity sends *signals ζ*_*i*_ to other entities, this signals reflect the *state* of the source entity but
they also depend on the values of a set of *parameters* 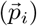; they are then propagated and modified by a non-local operator (**Φ**) to the sink 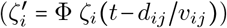. Once in the sink, the signal influences
4. A *functional interaction* is the net influence that a source entity excerts upon a sink entity. Therefore it has three components: i) the transformation from source state to the emitted signal 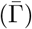, ii) the transport of the signal to the sink (Φ), during which the signal is modified, and iii) the transformation at the sink of the information carried by the received signal (Γ). A functional interaction ocurring at a given space-scale *s*_*l*_) for a given physiological function is therefore non-symetric and non-local (i.e, a structural discontinuity exists at that scale, the physiological mechanisms transporting the interaction information occur at lower space-scales and are different from those ocurring at *s*_*l*_).
5. At each space-scale *s*_*l*_) and at each time-scale (*τ*_*u*_), a physiological a function 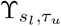 is observed; it is represented by a regulatory network 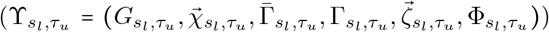, whose role is to maintain a given set of physiological variables within a certain (homeostatic) range even under perturbations, with responses ocurring as fast as possible within its characteristic time-scale. The regulatory network is composed of functional units and functional interactions among them (see figure 1).
6. Slower systems sense the mean activity of systems acting at faster time-scales and set the values of their parameters accordingly in order to maintain a “safe” or homeostatic regime.

**Figure 1:**
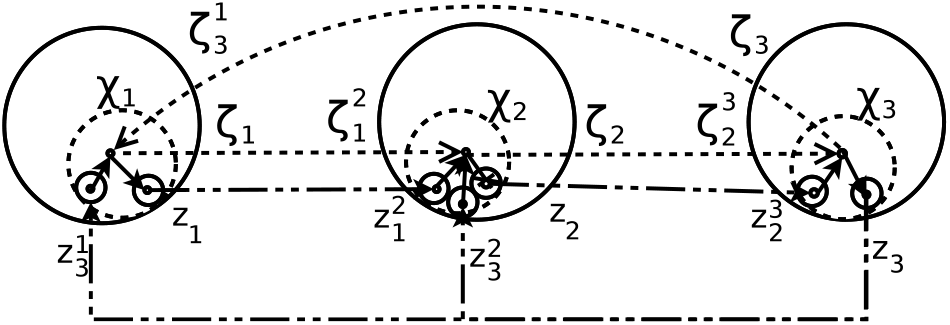
Simplified diagram of the hierarchical physiological system concept. A physiological system is represented by a multiscale regulatory network. At the higher space-scale level *l* = 1), a signal *ζ*_*i*_ representing the state *χ*_*i*_ of an entity (source) (e.g, excitatory) is sent to other entities (sinks), the signal received by a sink entity is nonlocal, has been transformed and possibly modified by other signals 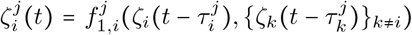. The high level signal can not travel through the physical space and must be first transformed internally at the source entity, going down in the scale hierarchy where an internal regulatory network processes it into signal 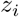 at the lowest level *l* = 0) (it might go down several scales until reaching the lowest level of description, here only two levels are depicted) 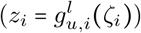 Signal 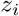 travels through a physical medium at the lowest scale that transforms it into 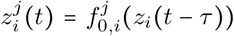 as it reaches the sink. The low scale signal 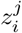 received by the sink is again transformed locally at the sink by a low scale regulatory network, where it may interact with other incomming signals, and it is promoted to a high level into an internal signal that integrates the information of several incomming sources 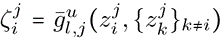. The state of the sink entity changes depending on set of internal signals 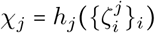.

**Figure 2:**
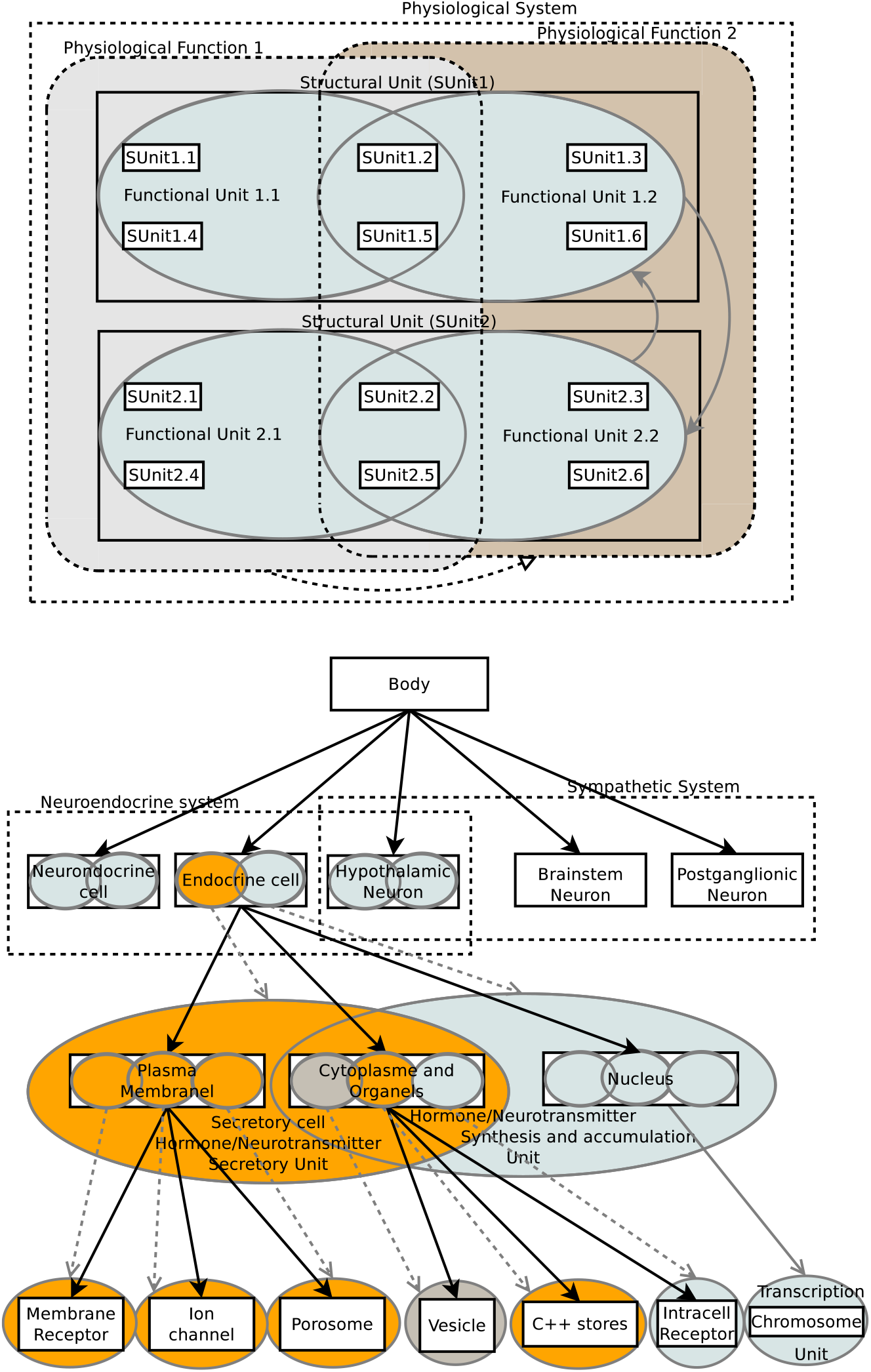
Relationship between structural and functional units

#### 2.1.1 Architecture (structural and functional organization)

- Structural unit (SUnit,): it is the basic structural element, living at a spatial-scale *s*_*l*_ (for *l* ∈ *L*, and *L* being the set of levels), it contains both a set of structural sub-Sunits, living at *s*_*l*−1_, and a set of Funits living at *s*_*l*_; the set of sub-Sunits may be void (in which case the Sunit is elementary) but the set of FUnits is not void. The two subsets are related as explained below.
- Functional unit (FUnit): is the elementary block of a physiological system; it lives at a spatial-scale *s*_*l*_ and a time-scale (*τ*_*u*_); it has a set of state variables, a set of parameters, and is capable of action, either in response to intrinsic (internal forces) or to external forces or both. It has one or multiple output ports to which it sends one or multiple signals. It may have one or more inputs from which it reads the information of incoming signals. It has an action mechanism giving the dynamics that determine the evolution the values of the state variables. It may contain a subset of functional units living at the same time-scale and at lower space scales.
- Elementary unit (EUnit): is the simplest structural unit that contains a single FUnit and no sub-SUnits.
- Each FUnit has a set of input variables, a set of state variables, and a set of parameters. In addition, it has a mechanism of state evolution; this mechanism may be implemented either by a sub-FUnit or directly by the FUnit using a given algorithm.
- There is a mapping between the set of FUnit inputs and a set of the inputs of their sub-FUnits, and another one for their set of outputs.
- For the set of sub-FUnits of a given FUnit (thus, working at the same time and space scales), there is a mapping between the their outputs and their inputs.
- There is a mapping between outputs of FUnits working at slow time-scales and the parameters of those working at faster time-scales.
- SUnits are simply containers of FUnits and other sub-SUnits
- Propagation operators are implemented as EUnits.

## 3 Results

### 3.1 Modelling the HPA system

The description of the basic molecular mechanisms involved in the function of the HPA system and the mathematical formulations are presented in the Appendix (A.1).

#### 3.1.1 Functional organization

The crucial and difficult point of the approach is the definition of the functional organization; physiological functions, together with the functional interations and functional units involved in them must be identified, as well as the signaling nature used by each functional interaction.

**Time scales and temporal behaviors:** At least four behaviors at different time-scales have been observed in the HPA system:

1. **fast** changes in ACTH concentrations occuring in the 10^1^ min scale (ultradian pulses or oscillations with 0.5 ~ 1 hour intervals[87, 88]),
2. **very fast** irregular fluctuations (mean interpeak intervals ~ 5 min) have been observed when measuring at a high temporal resolution (Δ*t* = 20 ~ 60 *sec*) in both horses [5] and primates [20].
3. **slow** changes in the ~ hour scale (circadian oscillations lasting ~ 24 hours, and
4. **very slow**, long lasting changes in the days scale under chronic stress or other sustained perturbations.

CRH concentration temporal profiles in pituitary venous blood are not well known in humans but there’s evidence that they are similar to those seen in horses, which show very small fluctuations and low concentrations (4 to 20 pmol/liter, higher concentrations have been observed in other species but measures may be flawed by stress), with only ~ 3 pulses/hour, compared to ACTH and AVP that show ~ 10 pulses/hour [5] (see figure 3). CRH basal concentrations are largely uncorrelated with those of ACTH, while AVP and ACTH show important temporal correlations (see figure 3); AVP seems therefore to be the main drive for high frequency ACTH secretion in horses and probably in humans in basal conditions while CRH seem to be the main activator during acute stress, its activity being strongly repressed by cortisol in basal conditions (but probably being sufficient to facilitate AVP action) [5]. Glucocorticoids negatively regulate HPA activity at multiple points with all CRH, ACTH and AVP levels being modified, but not rhytmicity [5].

**Figure 3:**
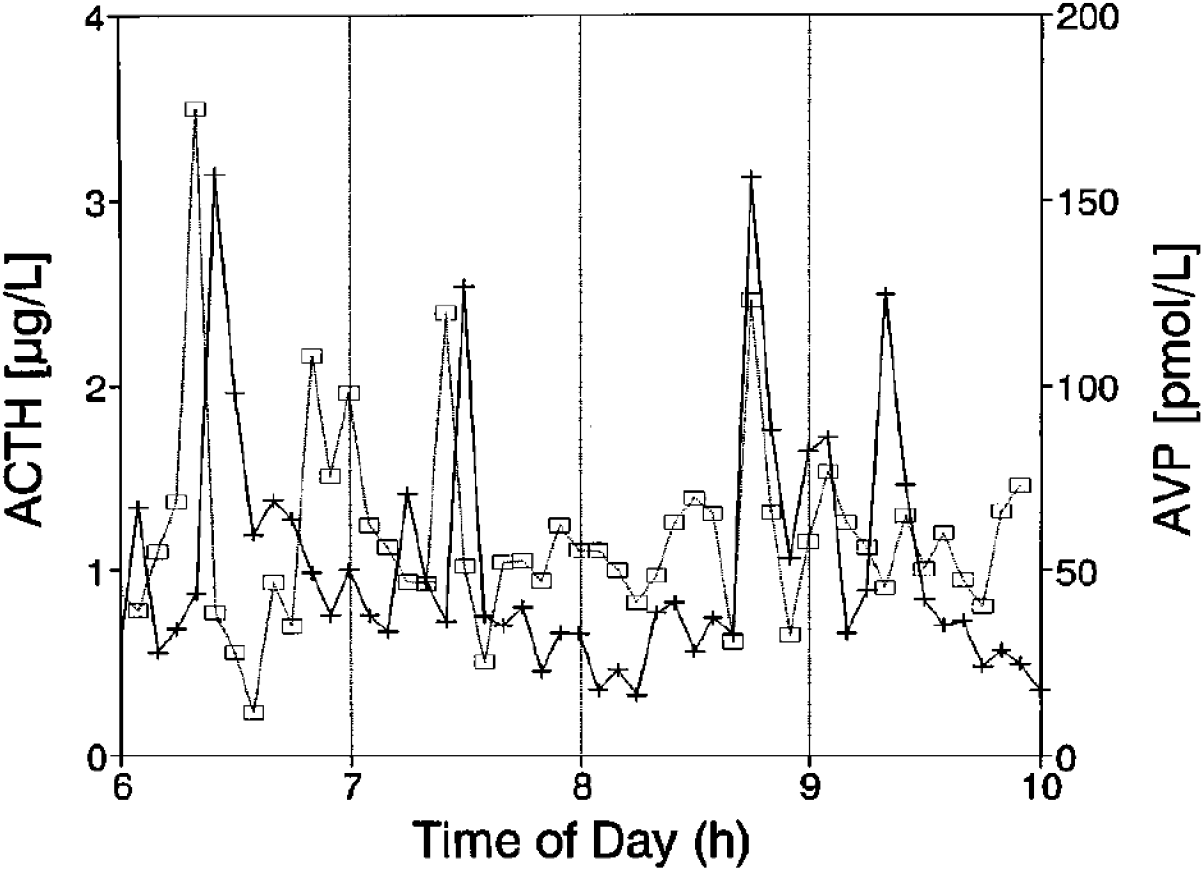
Pituitary venous concentrations of ACTH (plus) and AVP (open squares) in a typical horse bled at 5-min intervals. Redrawn from Redekopp *et al.* (161)

**Figure 4:**
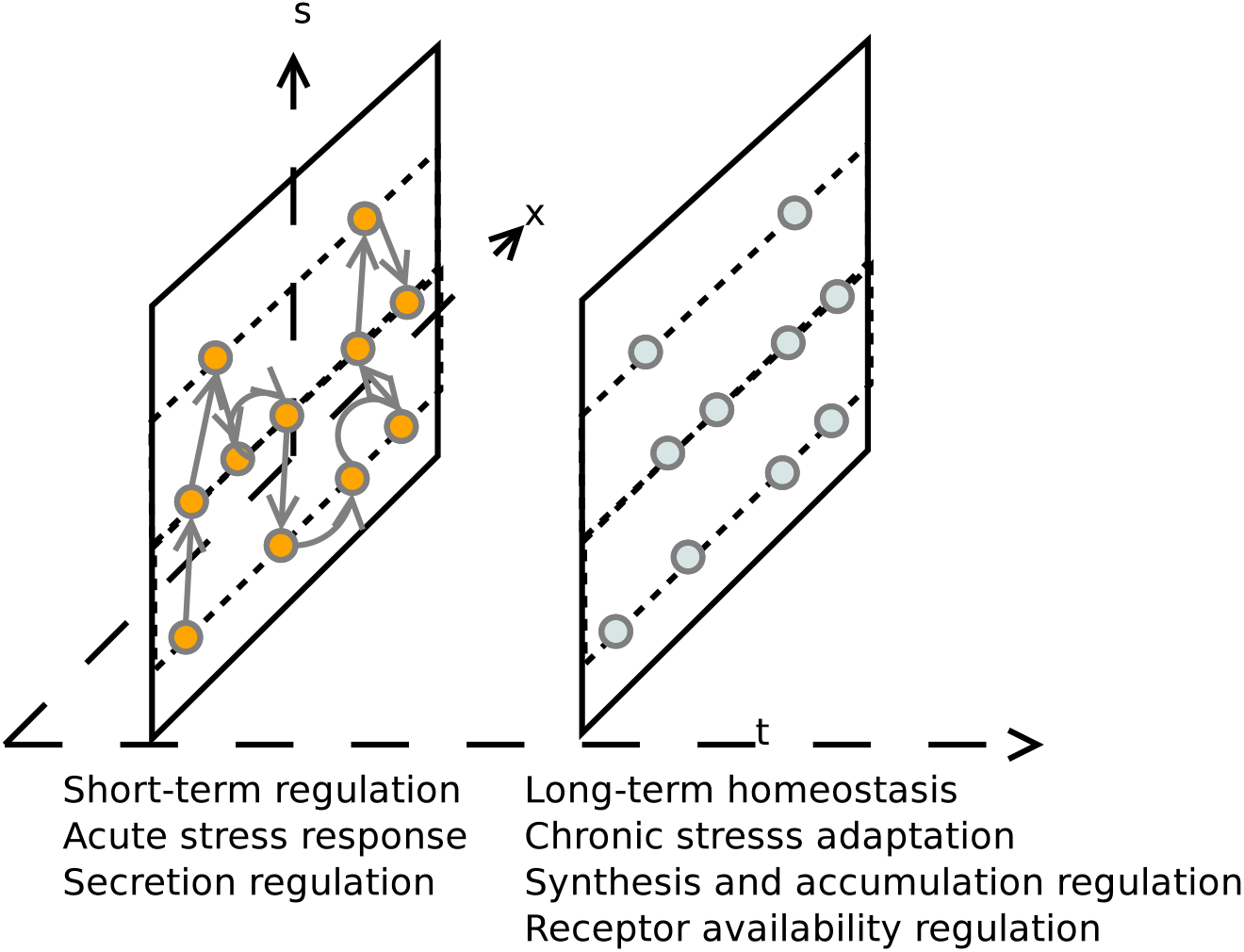
Physiological functions representation; functions that are upper in the functional hierarchy (masters), regulate functions down in the hierarchy (slaves) by setting their parameters; master functions may or may not ‘sense’ the activity of slave functions in order to regulate their activity (or equivalently, slave functions may or may not send information to the master function)

**Space scales:** We focus mainly on three scales of description:

- the scale of cell populations
- the scale of intracellular organelles
- the molecular scale

**Physiological functions:** given our interest in the analysis of clinical data of blood hormone concentrations, we focus on temporal behaviors that may be observed with these kind of data, ocurring from minutes to several hours. According to the time scales involved, there seems to be at least two main physiological functions:

1. Fast control of hormones concentrations mediated by the fast regulation of secretion rates.
2. Slower control of hormone secretion rates mediated by hormone synthesis and accumulation.

Circadian modulations may also be considered as a function of some HPA cells subpopulations given that some endocrine populations exhibit intrinsic circadian oscillations even in the abscence of external circadian drive from the SCN. However, most circadian drive comes from the SCN and modulates HPA circadian behavior (ref).

These two functions seem to be composed each of at least two subfunctions:

1. Fast control:

- Acute responses to stress (physical, emotional, immunological): mediated mainly by CRH and regulated by immediate cortisol feedback mediated by glucocorticoid receptors (GR) [106]; ACTH concentration is rapidly controlled by changes in the secretion rate. The cell uses fast internal signaling mechanisms to control secretion at its borders, depending on the internal cell parameters and on the incoming stimuli, but also on the extracellular milieu state in the vicinity of the cell. The amount of hormone ready-to-release is a parameter of this system that is set by the slow control homeostatic system. Fast cortisol feedback only reduces stress-stimulated but not basal ACTH [91].
- Short-term modulation of basal concentrations in response to osmolality and circadian conditions, mediated by AVP, and regulated by cortisol by delayed cortisol feedback acting on ACTH secretion.
2. Slow control:

- Homeostasis of long-term mean basal hormone concentrations: this is a slow process controlled mainly at the level of synthesis and storage by genomic mechanisms [91], its regulation by steroids is mainly mediated by mineralcorticoid receptor (MR) activation [106]. An additional control is excerted by the periphery (excretion, degradation) and by extracellular storage (blood protein-binding).
- Adaptation to chronic stress: CRH stimulation is inhibited during chronic stress [105] whereas AVP stimulation and response, regulated by AVP receptors (VR1b) expression [1, 3] are increased [105]

#### 3.1.2 Functional architecture of the HPA system

Known evidence on interaction links and functional units functions involved in the HPA system dynamics is summarized in the figure 5.

**Figure 5:**
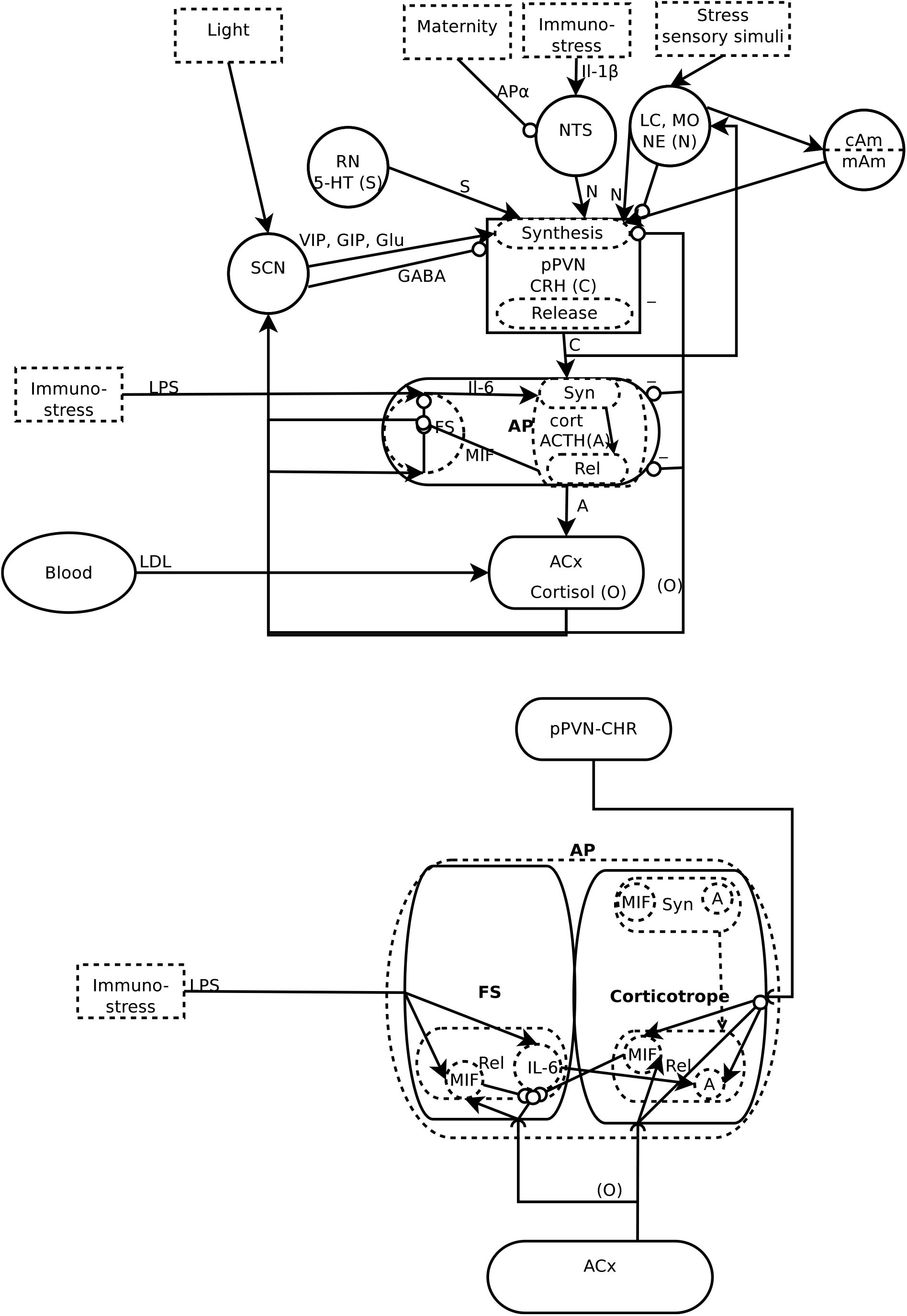
Evidence on functional interaction links of the HPA system and stress related inputs

#### 3.1.3 Structural and functional units

The basic structural unit at the higher scale of interest is the homogeneous cell population. Cell populations can be either neural, neuro-endocrine or endocrine. Every neuron is indeed a neuro-endocrine cell, because they use chemical signaling (secreting neurotransmitters, hormones, cytokines, etc), and endocrine cells show membrane excitability and electrical activities just as neurons do. There is other kind of signaling between neuroendocrine cells, the electrical gap-junction which transfers information directly from cell to cell without the need of neurotransmitters; however, this kind of signaling is internal to the cell population, allowing cell fast communication and synchronization across cell populations.

Even if the molecular mechanisms underlying neuroendocrine signal processing vary among different cell populations, a stereotypical basic internal functional process exist. First, incomming signals are transformed and amplified internally by different signaling pathways then internal signals trigger fast and slow responses; fast responses control the immediate amount of secretion, which depends on hormone/neurotransmitter availability; slow responses control the parameters of the fast process, such as hormone availability or sensitivity to incoming signals.

The actual mechanisms used by a cell depends on the specific type of cell, on incomming signals and on other factors. Some hormones are stored on vesicles (slow process controled) and released upon ocurrence of internal events or in response to external signals (fast process). Other hormones (e.g. steroids), freely diffuse, but the synthesis rate is controled by both fast and slow process.

**pPVN-CRH:** There are multiple populations of hypothalamic parvocellular CRH producing cells; some of them send their axons to the median emminence, but others project to other brain regions, such as limbic and brain stem regions. These neurons are regulated by neural control mediated by many neurotransmitters and by hormone feedback (both short and long range, i.e. from pituitary and other endocrine glands). Regulation by emotional stress mediated by cholinergic and noradrenergic neurotransmitters, but also by other peptides and endorphines [138].

*Synthesis:*

- Norepinephrine (NE) stimulates fast CRH gene transcription (upregulated at 15 min time-point after a microinjection in the PVN: 50*nmol*/100 *nL* over 10 s) and returned to basal levels by 120 min [72].
- GC regulation: Negative regulation of CRH concentration by glucocorticoids has been observed *in vivo* with great precision in horses [5], and there’s evidence of similar phenomena in other species. Upon disruption of cortisol synthesis, cortisol concentrations diminish slowly *τ* ~ 1*hr*) CRH concentrations rises abruptly (and probably in stages) after more than 1 hour of delay [5]; while the authors suggest a rate-sensitive feedback, the observed profiles suggest that a threshold (and probably multiple threshold) mechanism, relative to a set point, is more plausible. Constant infusion of a relatively small dose of 20 micrograms dexamethasone (DEX) did not reduce the hypothalamic CRF content; in contrast, the infusion of 202 and 504 micrograms DEX over 6 hr significantly reduced the hypothalamic CRF content [142]. These two evidences suggest a threshold mechanism for CRH synthesis regulation by GC.

**pPVN-AVP:** Negative glucocorticoid regulation of pituitary AVP secretion has been observed precisely in horses, just as with CRH; upon slow *τ* ~ 1*hr*) cortisol depletion after disruption of synthesis, AVP amplitude increases abruptly, but pulses frequency don’t [5]. Experiments where glucocorticoid flu-cutations have been prevented during NE stimulation of the pPVN suggest that AVP synthesis regulation by glucocorticoids is much stronger than that of CRH [72, 80]. NE injection caused an approximately 3-fold increase in AVP hnRNA levels by the 15 min time-point when cortisol synthesis is eliminated [72].

**Corticotrophs:** the corticotroph cell structural unit is composed of two main FUnits, the secretion FUnit and the synthesis and adaptation FUnit (see figure 9). The architecture of the secretion unit is depicted in figure 6.

**Figure 6:**
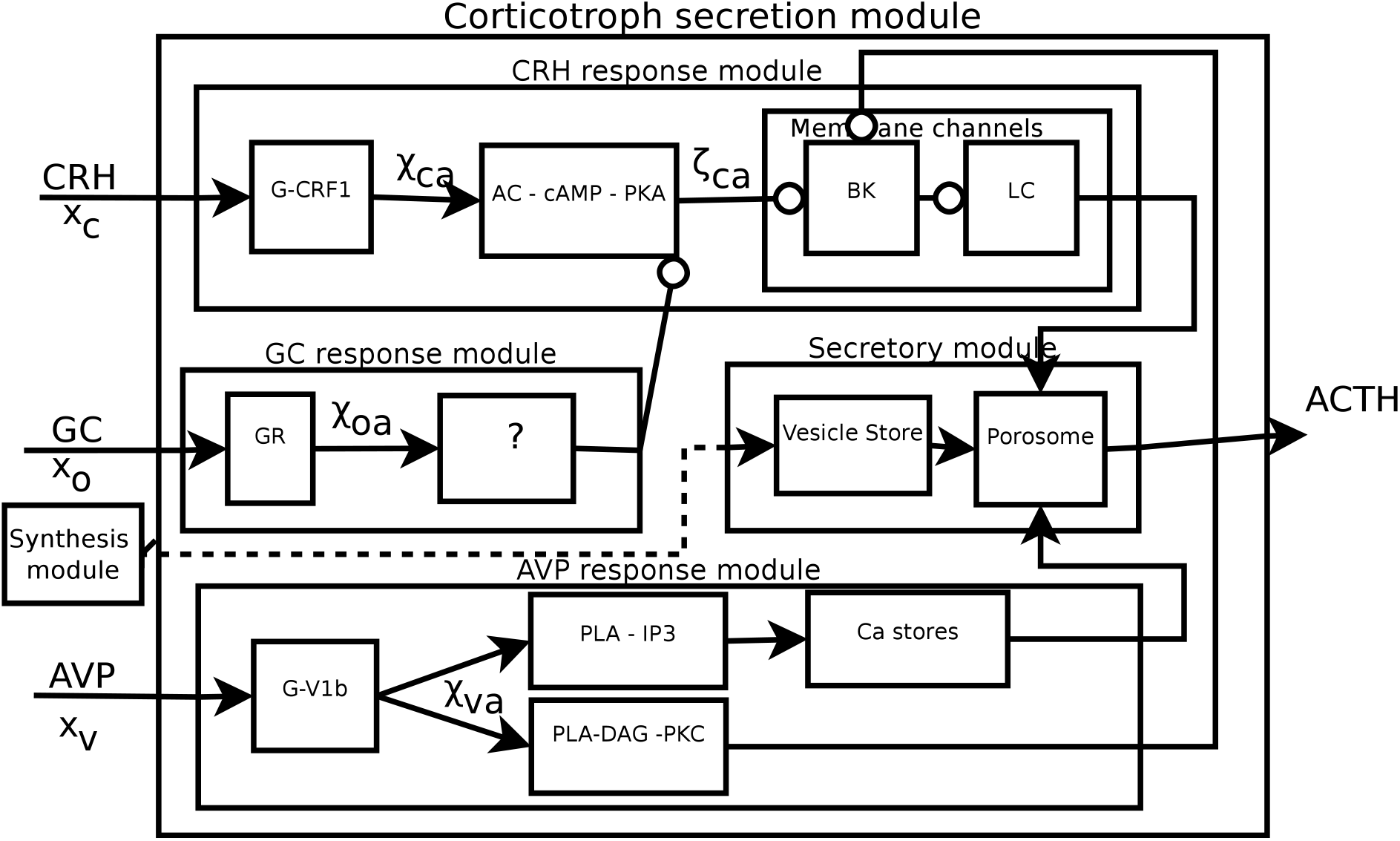
Corticotroph secretion functional unit (module). The number of available vesicles is a parameter of the system that is controlled by the synthesis functional unit (module)

*Global behavior:* in vivo measures of ACTH stimulatory and regulatory signals is very difficult in most animals and in humans. It has been shown (in horses) that stimulatory AVP and output ACTH concentrations are strongly correlated, both under hypertonic saline stimulus and in normal conditions, while CRH is largely uncorrelated with both AVP and ACTH concentrations in this conditions [5]. AVP and CRH elicit ACTH secretion using different pathways and produce different changes in ACTH concentration, CRH elicit graded responses (changes in hormone secretion per cell) while AVP produce non-graded ones (changes in number of secreting cells) [19]. While CRH seems to mediate ACTH secretion in response to acute stress, AVP seems to mediate adaptation to chronic stress (probably mediating proliferative responses in the pituitary) [1]. CRH stimulates secretion of ACTH by a factor of 8-fold in rat pituitary corticotrophs [174] with half-maximum effective concentrations (EC50) ≈ 3 × 10^−10^*M*. The main regulator of ACTH secretion is GC, which seems to mainly regulate ACTH release [142] by genomic and nongenomic pathways; while genomic pathways are relatively well known, nongenomic ones are much less known and more controversial [66].

**Secretory FUnit:** (figures 6, 7 and 9(left)). *CRH and AVP secretion stimulation:* CRH uses mainly the AC-cAMP-PKA -[*Ca*^++^]e ^1^ inux pathway, while AVP uses mainly the PLC - PIP2 - IP3 - [*Ca*^++^]_*s*_ ^2^ release patwhay, to increase the [*Ca*^++^]_*i*_ signal that stimulates secretion. However, it must be noted that recent results have suggested that both pathways are not completely independent, CRH can also stimulate *Ca* _*s*_ release via stimulation of both PKA and PKC kinases activity [45, 44] and PKC has been shown to influence CRF1 function [114]. In the sake of parsimony, we first assume independence of the pathways taking into account the best known phenomena and without taking into account all the complexity of the secretory network. The main reactions in response to CRH stimulus are shown in figure 7.

**Figure 7:**
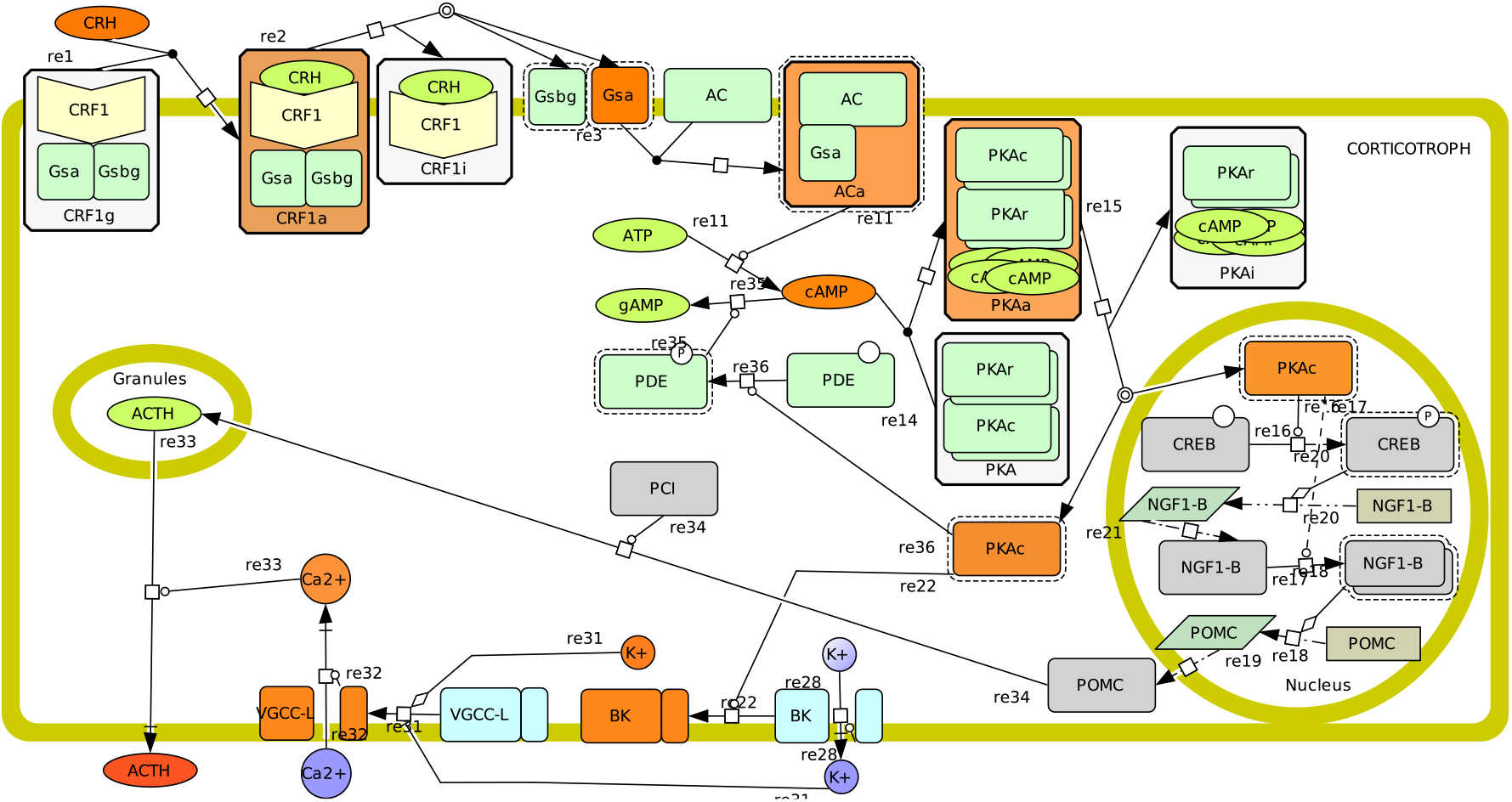
SBML notation diagram of reactions involved in the CRH control of ACTH secretion, a simplified view of the synthesis module is also depicted; reverse reactions are not shown for clarity of the signaling direction.

In vitro inhibition by corticosterone (in rats) of CRH-induced ACTH release has been observed in cultured pituitary explant fragments; it occurs within < 20 min and is functional in the presence of cycloheximide, suggesting a nongenomic mechanism of action [50]. There are reports that fast increases in cortisol concentration rapidly inhibit ACTH secretion apparently using a rate-sensitive mechanism [91, 106] (however, it is not clear how such rate-sensitive mechanism would work or even if it is really at work). Glucocorticoids excert a multimodal regulation on CRH triggered secretion, with varying time delays *min ~ hour* scale) and at different sites: i) it acts relatively fast at the input level inhibiting CRH binding to CRF1 receptors (15 ~ 60 *min*) (unknown mechanism) [29]; ii) it can act rapidly ~ 15*min*, at the output, involving FS cells and paracrine signaling [152] and probably inhibiting vesicle excretion mechanisms [17, 85], iii) it can act on membrane potential and action potential firing, supressing *K*^+^ channels inhibition by the CRH pathway. For this last phenomenon, indirect (probably genomic) mechanims acting in ~ 1*hour* have been proposed [149, 161, 162] but also a fast (~ *min*) and direct mechanism [79] (however it is not clear if the second one is actually functional *in vivo* because it requires high GC concentrations *EC*_50_ ≈ 21*μM* [79]). An hypothetical alternative pathway may involve the regulation of Ca^++^ feedback [9], but it is less well documented.

- *Input signals reception and transduction:* CRH,AVP and CORT bind to receptors and activate intracellular signaling pathways. CRH binds the CRF1 GPCR at the cell membrane with high effective affinity to its receptor thanks to a two-domain affinity “trap” mechanism [77], facilitating coupling to the receptor and activation of a Gs protein [14]. CRH dependent Gs activation [14] and cAMP acummulation [78] at equilibrium (measured *in vitro* during 30 min incubation for cAMP [78] and 120 min for Gs [14]) was described by a sigmoid concentration-signaling function. A kinetic model has been proposed in [78] assuming constant concentration of Receptor. However, this is a trimeric reaction and it has been suggested that glucocorticoid modulation of the receptor avail-ability could an important variable of the CRH response [29] and Gs protein availability could also be one. In addition, there is *in vitro* evidence (in rat corticotrophs) that CRH-triggered secretion last for 10~15 min after CRH washout [93], which may indicate that the strong effective affinity of CRH and its GPCR results in a long dissociation time, as it has been suggested for cortisol and the MR receptor [106], and therefore a long-lasting activation of Gs proteins by the same CRH-CRF1 complex. Therefore, we assume that i) CRF1 concentrations may be quickly modulated by other signals, and ii) that responses to CRH short pulses rise fast (~1 min) and decay more slowly (~10 min) independent of other signals. *CRH stimulus:* CRH binds the CRF1 receptor at the cell membrane which then activates a G protein. The equilibrium expression for the pituitary CRF1 response is then given by (37):

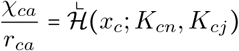

where *χ*_*ca*_ is the transducted CRH signal (c) in the adenohypophysis (a) (the activated CRH-CRF1 complex concentration), *r*_*ca*_ is the total CRF1 membrane concentration in each corticotrophe, *K*_*cn*_ are the affinity constants for CRH binding to the N domain of CRF1, *K*_*cj*_ is the isomerization constant for the J domain of CRF1; *K*_*c*_ = *K*_*cn*_ (1 + *K*_*cj*_) is the macroaffinity contant for CRH binding to CRF1. It has been shown that if CRF1 is not coupled to a G protein both affinities are very weak, while for the G-coupled receptor (G-CRF1) they strongly increase [78] (see A.2). Therefore, we may neglect the uncoupled receptor state and we may assume that *K*_*cn*_ << *K*_*cj*_ and *K*_*c*_ ≈ *K*_*cn*_*K*_*cj*_ for the coupled state, which gives a simplified expression for the G-CRF1 receptor transfer function at equilibrium for CRH:

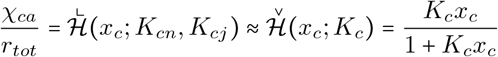

and the dynamics of the *G*_*sα*_ signal response to CRH may therefore be approximated by:

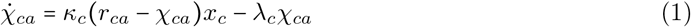 There are several sources of information concerning the dynamics of ligand binding to the receptor [93, 174, 14, 78] and of the whole receptor system including G-protein activation [14], from where parameter estimates for the equilibrium parameters of the whole receptor system (*K*_*c*_) can be drawn. However, no data was found from which reliable estimates of the kinetic constants (*κ*_*c*_,*λ*_*c*_) may be obtained; educated guessed are given in table 4, see A.2.3 for estimation details. Using *κ* ≈ 0,95(nM *min*)^−1^ and λ_c_ ≈ 0.2/*min* and for a saturating concentration step (~ 10 *nM* [93]) the rise time of the response is very fast (τ ≈ 0.1, see A.2), which is coherent with the observation that the onset of responses to CRH occurs very fast (< 1*min* [93]) and that CRH is a potent secretagogue. *cAMP - PKA signaling:* i) the CRH-activated, GTP bound Gs protein binds and activates the Adenylate Cyclase (AC); AC transforms ATP on cAMP, which activates the PKA enzyme, which finally acts on the activity of membrane ion channels and on other targets. We assume that the dynamics of this signaling pathway are given by equations (42) and (41) (see A.1.2):

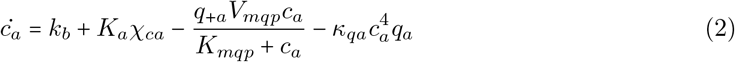

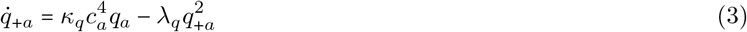

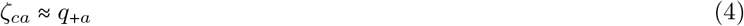

where *q*_*+a*_ and *q*_*a*_ are the active and inactive PKA, respectively;, *k*_*b*_ represents the basal AC (enzyme) activity;; *V*_*mqp*_, *K*_*mqp*_ are the Michaelis-Menten constants of the PKA-PDE negative feedback; *χ*_*ca*_ is the input signal (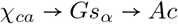 is neglected, assuming fast and unit gain trans-transfer, see A.1.2) and thus *K*_*a*_χ_*ca*_ is the maximum rate of cAMP synthesis; and *c*_*a*_ = [*cAMP*] is the cAMP intermediate signal. The effective intracellular signal of this pathway (*ζ*_*ca*_) is assumed to be the active PKA, which excerts then fast effects on electrophysiological activities in a few minutes. *AVP stimulus:* the pituitary AVP receptor (V1b) is a GPCR that signals through the Gq protein. Assuming the same simplifications as for G-CRF1 (1) the dynamics are given by:

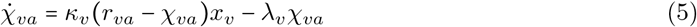 Experimental values for *κ*_*v*_ and *λ*_*v*_ are given in A.2, table 4, taken from [61]. *PLC-IP3 mediated* [*Ca*^++^]_*s*_ *release:* the activated V1b receptor Gq protein signal activates PLC, which++synthetizes IP3, which then stimulates an increase on [*Ca*^++^]_*i*_ by release of intracellular stores ([*Ca*^++^]_*s*_) [164, 166]. *GC signal transduction:* at the pituitary it occurs via both GR and MR, GR seem to detect peak glucocorticoid concentrations, either of basal pulses or induced by acute stress, while MR which seem to be active most of the time and probably carry information on mean basal concentrations [106]. Glucocorticoids bind with higher affinity to the MR than to the GR receptor. There are at least two different kind of transduction for GC : i) binding and activation of intracellular receptors followed by nuclear translocation → DNA binding → transcription regulation, mediated by intracellular GR and MR, or ii) binding and activation of GR membrane receptors followed by a nongenomic signaling pathway probably involving protein kinase C (PKC) and MAPK kinase [152]. Even if this last mechanism was observed in FS cells, it might also work in corticotrophes. We may thus assume that fast GC inhibition of the CRH input depends on a reaction chain dependent on this last mechanism, while actions on ACTH synthesis depend on the slower transcriptional GR and MR pathways. *GR transduction:* membrane GR features are not well known, therefore we may assume that it behaves similar to the intracellular GR, which has relatively fast half time dissociation time *τ*_1/2_ ≈ 5*min*. Therefore, the cortisol transduced signal is given by:

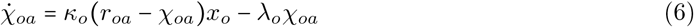

with *λ*_o_ ≈ *ln*(2)/5*min* ≈ 0.14/*min* and assuming a constant number of available GR in each corticotrophe *r*_*oa*_. *Desensitization/enhancement of GPCR function:* homologous desensitization was found to be rapid for V1b (in ovine corticotrophes it was complete within 5 to 10 min of the onset of AVP treatment, for concentrations of (5nM); this is within the range of concentrations and durations of VP pulses seen in sheep portal blood during acute stress) and readily reversible (resensitization was complete 40 min after the end of AVP treatment) [115, 69], whereas for CRF1 is much slower (requiring longer than 25 min with a CRH concentration of 1 nM) [115, 114]. One possible pathway for homologous or heterologous desensitization/enhancement of CRF1 function involves PKC activity; PKC modifies CRF1 function by phosphorylation, but the kind of influence depends on the variant of the CRF1 receptor; however, the main function of PKC with respect to CRH responses seems to be the enhancement of AC-cAMP pathway activity and may partly explain the amplification of AVP on CRH responses [114]. For V1b it depends on receptor internalization and resensitization depends upon PP2B-mediated receptor dephosphorylation. *Membrane depolarization:* A model of action potential generation and [*Ca*^++^]_*i*_ signaling was proposed by [97, 98, 150] that includes L and T type VGCC currents as well as two (*K*^+^) ionic currents including BK type; we base our model on this work. However, as we are not interested in detailed dynamics ocurring at the msec scale but at slower time-scales (mean membrane voltage and firing rate changes ocurring during minutes), we simplify the model accordingly (see A.3 for details and for the precise currents formalizations).
- CRH-AVP-CORT intra- et intercellular interactions: CRH and AVP both increase the binding of each other [29, 30]. GC inhibit CRH binding to cells but don’t interfere with AVP binding. The precise mechanisms involved in AVP and CORT influences on CRH binding are unknown but it has been shown that CORT action in CRH can be very fast and thus involves non-genomic mechanisms [50]. Slower but relatively fast (*τ* ~ 30*min*) interactions occur at later stages in the signaling pathway: CRH-triggered depolarization, *Ca*^++^ entry and subsequent ACTH secretion, are inhibited by CORT and potentiated by AVP; apparently these interactions occur at the level of the voltage and *Ca*^++^ dependent BK (big *K*^+^ conductance) channels, which are inhibited by both PKA and PKC and this inhibition is itself inhibited by CORT [156]. *GC fast regulation of CRH input:* Fast regulation at the input has been shown to involve a strong decrease in the percentage of CRH-bound corticotrophes [29, 30]. Evidences that CRH-triggered secretion last 10~15 min after CRH washout in rat corticotrophes [93], that the CORT-induced ACTH release inhibition is observed within ~20 min [50], and that the percentage of CRH-bound cells is reduced by 50% in ~10 min after exposure to glucocorticoids [29, 30], suggest that CORT acts rapidly at the level of the CRF1 receptor. There are at least two possible mechanisms: i) interference with the CRF1 availability at the membrane [29], or by interfering with the GCRF1 function (inhibiting G-protein coupling and/or CRH binding). In both cases, we may imagine a simple competitive reaction that makes the CRF1 unavailable, modifying (1) into:

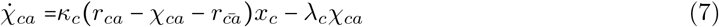

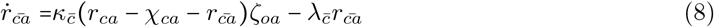

where 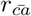 is the fraction of total GCRF1 (*r*_*ca*_) that is not available, and the kinetic parameters 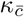 and 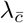 are unknown.
- *Electrical activities:* Corticotrophs exhibit both spontaneous large-amplitude spiking and pseudo-plateau bursting [156] and this electrical activity is modulated by several hormones, CRH [92, 93, 94] and AVP [34] being best known and most important. Action potential generation depends mostly on calcium currents through L-type *Ca*^++^ voltage-gated calcium channels (VGCC) (L for longlasting [62]) while firing rate is modulated by P-type ones [92]; *NA*^+^ currents have a negligible role [93, 94]. cAMP dependent PKA activation in response to CRH triggers a persistent membrane depolarization via a closure of background K channels[93, 165] [100] facilitating action potential firing and increasing membrane resistence [92]. The onset of depolarization and firing rate increase occurs rapidly, after a delay of about 45sec and its effects last over 10 to 15 min after CRH stimulus offset [93]. However, the CRH-induced firing rate increse is only partially dependant in membrane potential decrease, suggesting that other CRH dependent phenomena may account for the additional increase in firing rate [92]. In rat, the firing frequency increases to (~ 22*spikes/min*) as membrane potential diminishes compared to spontaneous frequency (∼ 12 spikes/min) (at ~ −50mV) or 0 in obtain a maximal stimulation of ACTH initial release rate [93], individual spikes last ≈ 1*sec* and have maximum amplitudes ≈ 20*mV* [92]. Membrane potential rises to about ~−50mV from the resting potential (~ −65mV) for quiescent cells or ~−55mV for spontaneous firing cells [92]. Application of 100 nM AVP slightly depolarized (≈ 5*mV*) [34] *Membrane depolarization:* there are several ionic channels expressed in the corticotroph membrane that control the membrane potential. We base our formulation on the one proposed by LeBeau et al. [97], who model action potential generation and [*Ca*^++^]_*i*_ signaling by including L and T type VGCC currents (*I*_*LC*_, *I*_*TC*_), DR and calcium regulated BK type (*K*^+^) currents, and later an inward rectifying *K* current important in the triggering of depolarization and spiking [150]. However, we retain only those that are essential for the setting of the mean membrane potential (L-VGCC, BK and DR *K*^+^)), as we are not interested in detailed dynamics ocurring at the msec scale but at slower time-scales (mean membrane voltage and firing rate changes ocurring during minutes), we simplify the model accordingly (see A.3 for details).

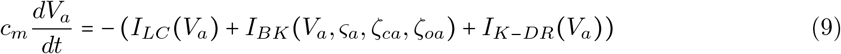 However, some important modifications must be done to LeBeau et al. model in order for the model to account for new knowledge on BK channel mechanisms. First, their assumption that PKA modifies the voltage dependence of the *I*_*LC*_ current by directly phosphorylating the channel [97, 98] is not supported by any experimental evidence; however, new alternative evidence exists showing that PKA directly inhibits the BK channel activity instead [148, 149, 26], therefore, we modify the equations accordingly (PKA is known to inhibit the e21(STREX) variant of the BK channel, which is expressed in the pituitary, and to activate others [26, 160], we may assume therefore that the e21(STREX) BK channel variant is the one mainly expressed in corticotrophs membrane (as has been shown to be the case in mouse cells [148]) and functionally involved in secretion stimulation by CRH). In addition, it must be noted, that recent reviews on the BK channel function show that [*Ca*^++^]_*i*_ act as an allosteric factor ^3^, facilitating the opening of the channels[37, 26, 96], rather than directly activating them. Moreover, the experimental voltage dependence of BK channels conductance follows a sigmoid response curve, with the G-V curve shifting right as [*Ca*^++^]_*i*_ increases, and reaching saturation at high voltages; on the contrary, LeBeau et al. model BK conductance formulation is expressed as the product of a constant and a Hill function of [*Ca*^++^]_*i*_. *GC regulation of membrane depolarization and firing rate:* a second point of GC regulation of secretion acts on voltage and *Ca*^++^ dependent *K*^+^ channels function. A first relatively fast mechanism involves the inhibition of PKA [149] and PKC [161, 162] and the phosphorylation of BK channels [156]; however, it involves mRNA and protein synthesis and therefore acts at lower speeds. The IC50 of dexamethasone to block CRF-induced ACTH release was ≈ 7.7 ± 1.9*nM* [149]. Therefore, we retain a simplifed LeBeau et al formulation for *I*_*LC*_, retaining its steady-state properties, while using an allosteric formulation for *I*_*BK*_ channels dependence on *Ca*^++^]_*i*_ and PKA, as well as GC signal inhibition of PKA action (see A.3 for details):

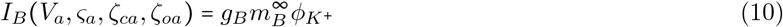

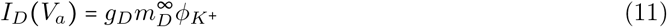

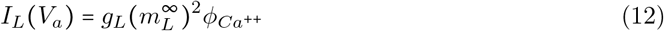

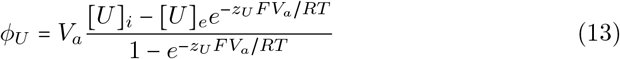

where *V*_*a*_ is the membrane potential; [*U*]_*i*_ and [*U*]_*e*_ are, respectively, the intracellular and extracellular ion concentrations for each channel; *ς*_*a*_ = [*Ca*^++^]_*i*_ is the intracellular calcium concentration; *m*_*x*_ are activation variables whose steady states are given by 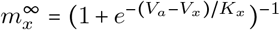, for *x* ∈ {*D*, *L*}, and 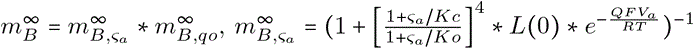 being the open channel probability dependent on *Ca*^2+^ [37] and 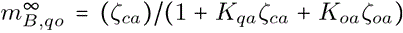 the open probability dependent on *ζ*_*ca*_ and *ζ*_*oa*_ (assuming independence of PKA and *Ca*^2+^ action. *Kc*, *Ko* are the closed PKA and for the GC signal (assuming a competitive mechanism for GC inhibition of PKA action). *L*(0) is the open-to-closed equilibrium constant in the absence of bound *Ca*^2+^ at 0 *mV* and Q is the equivalent gating charge associated with the closed-to-open conformational change [37]. *Firing rate:* We assume a nonlinear sigmoid function relating the decrease on membrane potential, due to the closing of background K^+^ channels, to the firing rate of VGCC-mediated *Ca*^++^ action potentials. We neglect other potential factors modulating the firing rate increase.

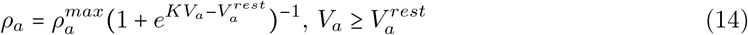 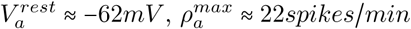
- *Intracellular calcium dynamics:* basal [*Ca*^++^]_*i*_ ≈ 60*nM* in quiescent cells, spontaneous [*Ca*^++^]_*i*_ waves depend mostly in L-type channels activity and reach maximum [*Ca*^++^]_*i*_ ≈ 300*nM* (in cultured rat cells) and mean values are ≈ 150*nM*; under a strong step CRH stimulation (20*nM*), the basal concentration suffers a persistent increase (even after CRH washout) up to ≈ 300*nM* in 5 min and ≈ 500*nM* in 10 min; similar results under 1*nM* (Bu)_2_cAMP [92]. [*Ca*^++^]_*i*_ waves show rising times 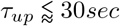 and show decreasing times of *τ*_*up*_ ~ 1*min* [92], which depend on action potentials elicited by L-type channels activity[92]. These waves result from the accumulation of small amplitude transients with 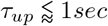 and decay times of several seconds [34], which are correlated with action potential spring and its frequencies (≈ 0.31 ± 0.22 Hz) [34] correspond to firing rate (≈ 0.36 Hz [92]), [*Ca*^++^]_*i*_ amplitudes being thus correlated to the firing rate [34]. Most *Ca*^++^ influx depends on action potentials but a small a mount on increase on basal levels upon CRH stimulation is independent of action potentials and depends on the CRH-induced membrane depolarization. Following [97] it can be expressed as:

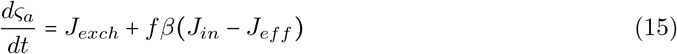

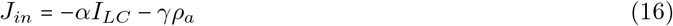

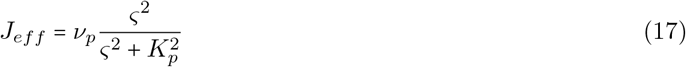

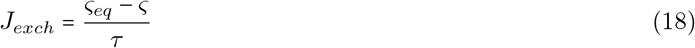

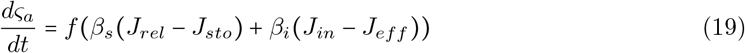

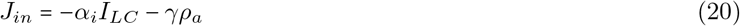

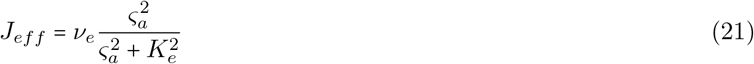

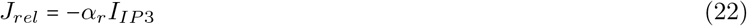

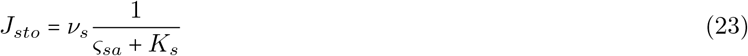 State variables: *ς*_*a*_, *ς*_*sa*_ intracellular and stored calcium, respectively; inputs: *I*_*LC*_, *ρ*_*a*_; parameters: *ς*_*eq*_, *τ*, *α*, *γ*, *ν*_*p*_, *K*_*p*_, *f*, *β*. Where *ς*_*eq*_ is the equilibrium [*Ca*^++^]_*i*_] value at rest, *f* is the cytoplasm buffering factor, *β*_*i*_ is the ratio of the cell membrane surface to cell volume and *β*_*s*_ is the ratio of the endoplasmatic reticulum membrane surface to cell volume. Compared to the expression in [97], additional release and storage *Ca*^2+^ currents through IP3R channels (*J*_*rel*_, *J*_*sto*_) have been added in order to take into account AVP action.
- *Secretion/Replenishement:* Similar to neurons and neuroendocrine cells, corticotrophs secretion is induced by an increase on intracellular *Ca*^++^([*Ca*^++^]_*i*_), caused partly by CRH-induced depolarization and action potential firing increase which activate L-type and other VGCC currents [93, 165], and partly by AVP triggered *Ca*^++^ release from intracellular stores [164, 34]. *In vitro* evidence depolarization shows that a 2 *Hz* (250 *msec* duration) train of depolarizations deplet the readily releasable pool (RRP) of granules in 5 sec (10 pulses), each pulse. Secretion can also occur via intracellular Ca^++^ release [165]. AVP induces intracellular Ca^++^ release producing a Ca^++^ ‘spike and plateau’, and NE elicits a rhythmic Ca^++^ release; CRH is the more efficacious despite the smaller Ca^++^ increase amplitude [165]. The peak Ca^++^ elicited by a train of 10 depolarizing voltage steps to +10 mV (250 ms duration; 2 Hz) ranged from 2·2 to 4·5 *μ*M. The rate of exocytosis increased with as Ca^++^ increase, at 2·4 *μ*M the rate of granules release was of approximately 63/sec .. *Secretion:* it is given by equations (47) and (49); ACTH release during exocytosis is highly efficient [127] so we may neglect partial secretion (*q*_*a*_ = *θ*_*a*_).

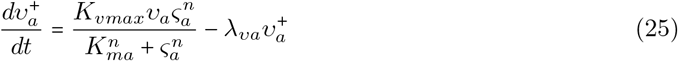

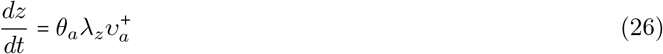 *Readily-releasable pool replenishement:* given by equations, and. We assume a strong basal synthesis of POMC that continuosly sets the replenishement rate to its maximum *K*_*smax*_ such that any change in replenishement is excerted by a slower genomic system changing the value of this parameter.

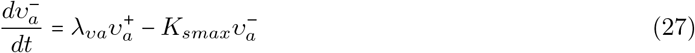

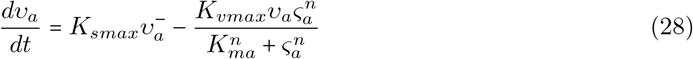

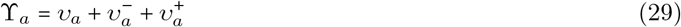 State variables: 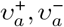, *v*_*a*_ (activated, inactivated and available hormone vesicles); input: *ς*_*a*_ (intracellular calcium) and *ξ*_*a*_ (synthetized POMC); parameters: *K*_*vmax*_, *K*_*ma*_, λ_*va*_, ϒ_*a*_ Putative parameter values estimated from (rat, *in vitro*) data in are *n* = 3, *K*_*ma*_ = 19 ± 2*μ*M (taken from figure 8 in [165]), *v*_*a*_ ≈ 4708 vesicles (see A.2 (values for human corticotrophs are not available).
- *GC regulation of ACTH exocytosis:* a third, non-genomic, mechanism for fast GC secretion regulation involving paracrine interactions and acting at the level of vesicle exocytosis has been proposed [152, 17, 84]. It follows an intracellular signaling pathway in FS cells, involving protein kinase C (PKC) and MAPK kinase [17], and phosphorylation and translocation of the protein Annexin A1 (ANXA1) to the plasma membrane of FS cells; subsequently it inhibits ACTH secretion on corticotrophes, probably mediated by members of G-protein coupled formyl peptide receptor family (FPR1/2) [152, 17, 84]. The GC-induced cellular exportation of ANXA1 in the rodent anterior pituitary gland parallels the onset of the steroid inhibition of ACTH secretion, both emerging within 15 min and reaching a maximum within 2 h

**Figure 8:**
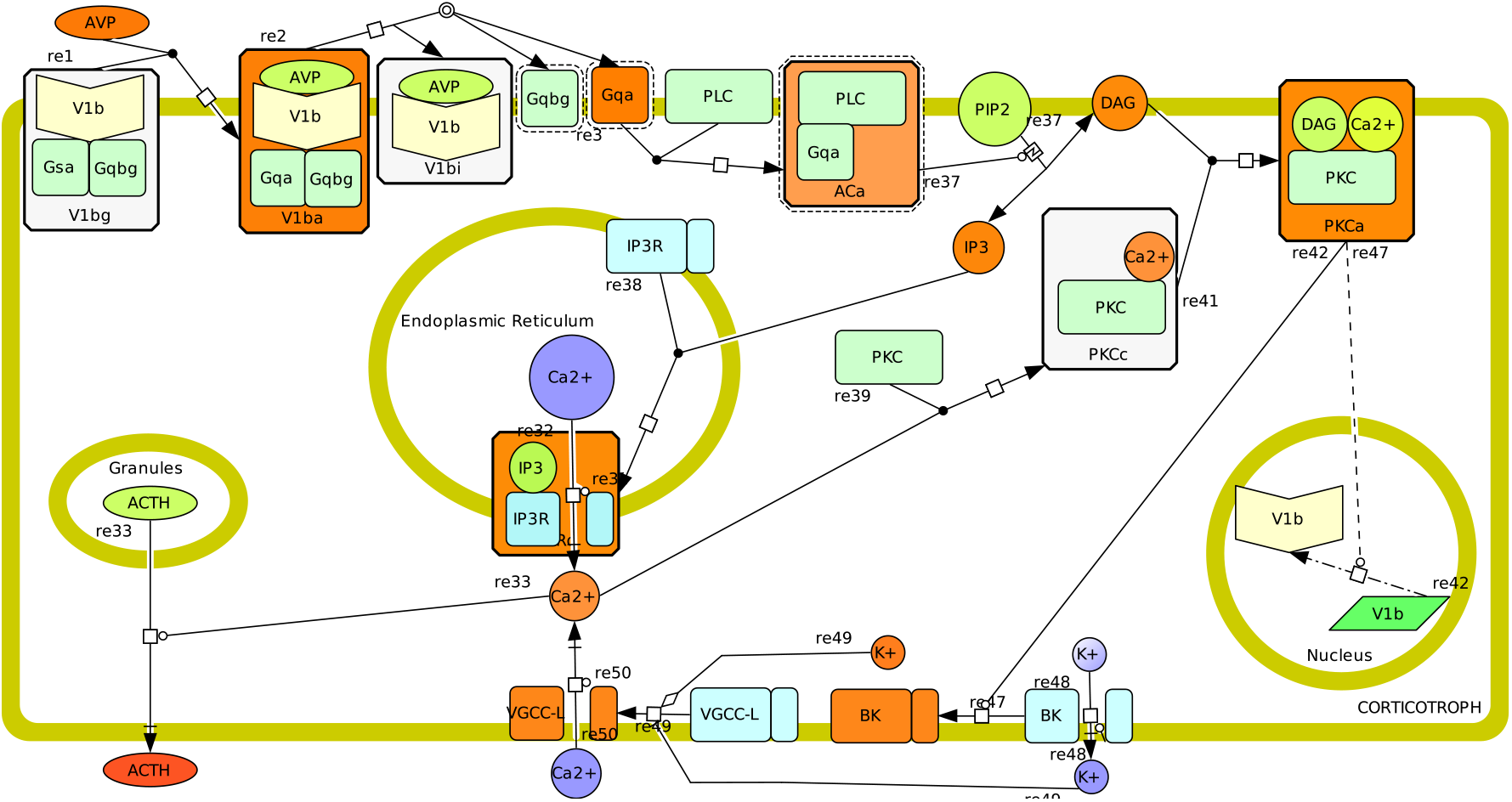
SBML notation diagram of reactions involved in the AVP control of ACTH secretion, reverse reactions are not shown for clarity of the signaling direction. AVP binds the V1b receptor activating a Gq protein, which itself activates binds to a PLC enzyme activating it. PLC cleaves a membrane phospholypid (PIP2) into two active molecules, DAG and IP3. IP3 binds to IP3R channels allowing *Ca*^2+^ release from intracellar stores. DAG activates protein kinase C (PKC) together with intracellular *Ca*^2+^. PKC targets multiple cellular processes, including extracellular *Ca*^2+^ inflow by inhibiting BK *K*^+^ channels and by indirectly activating V1b receptor transcription (not shown) and translation. Homologous desensitization of V1b receptors in response to AVP is fast and important, but its molecular mechanisms are unknown and thus it is not shown in the graph.

**Synthesis FUnit:** see figures 6 and 9(left)

**Figure 9:**
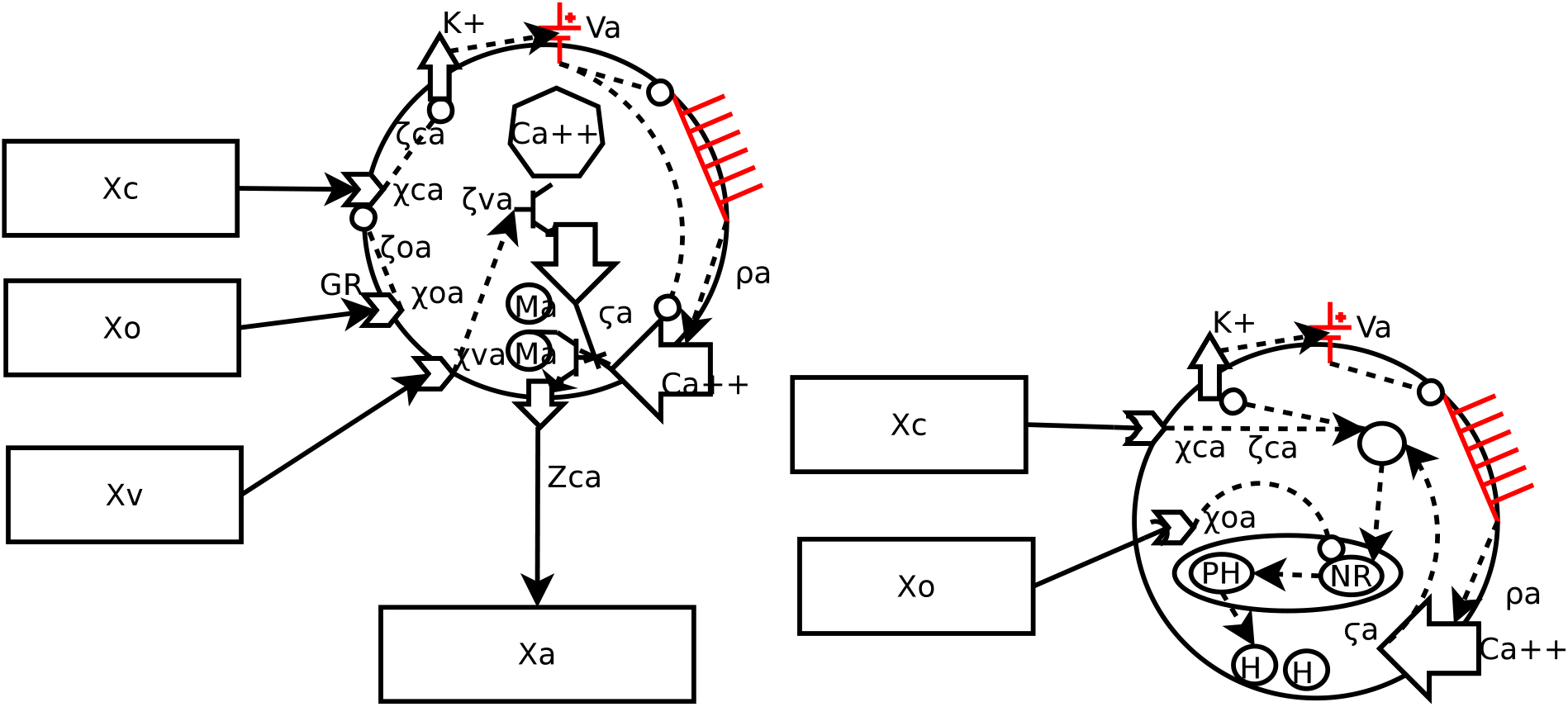
Left: secretion control in corticotroph. CRH excerts a fast and sustained control in response to acute stress by eliciting a sustained *Ca*^++^ inflow through depolarization and firing rate increase, whereas AVP elicits fast liberation of internal *Ca*^++^ stores in response to other signals. CRH signal reception and transdution is rapidly inhibited by glucocorticoids throught the GR receptor. Right: synthesis control. CRH elicits POMC transcription from which ACTH is then produced, while intracellular GR activated by cortisol goes to the nucleus and repress POMC transcription.

- *ACTH synthesis regulation:* Regulation of the ACTH precursor gene (POMC) is mainly transcriptional. The Ikaros (Ik1), Tpit and Pitx1 proteins that are present in corticotroph cells, bind to the POMC promoter and upregulate its expresssion, they are needed for normal POMC transcription as well as for corticotroph differentiation and population growing [52, 51]; Ik1 functions as a potentiator of transcription by remodeling the densely packaged chromatin environment of POMC, allowing activation by Tpit and coactivators [52]. Promoter elements are present in the POMC gene region that are responsive to stimulation by CRH and AVP [154]. Long-term chronic stimulation with CRH leads to its upregulation, while sustained exposure to glucocorticoids downregulates it. CRH upregulates POMC transcription indirectly through cAMP-PKA-dependent NGF1-B activation (potentiating NGF1-B dimerization and binding to NuRE promoter elements and recruitement of SRC coactivators, without *de novo* protein synthesis); NGF1-B activates transcription in concert with SRC [112]. Basal synthesis, independent of stimulation, is mainly dependant on Tpit/Pitx, while hormone responses are mainly mediated by the NuRE elements [122], but both elements participate to both kind of activities [122]. Both CRH and AVP have been reported to stimulate POMC transcription in ovine fetal corticotrophs (after long stimulation ≈ 92 *h*) [116, 123], AVP in adult cells [167], and CRH in atT20 mouse cells (observed at ≈ 72 *h*) [154] (nothing observed at 3h in [21]) in [21]) and rat (after a 30-min treatment with 0.5 nM CRF, POMC primary transcript levels were increased by 200-400%, 18-h incubation with 0.5 nM CRF increase cytoplasmic mRNA levels to about 140%, while AVP affect transcripts processing)[103]; however, while in one study only CRH seems capable of reducing GC inhibition of POMC regulation in mature cells according to [123], in other study only AVP is reported to increase POMC synthesis [167]. *GC regulation of ACTH synthesis:* experiments in horses show an upregulation of ACTH amplitudes (but not pulse frequencies) when cortisol concentrations diminish slowly after inhibition of synthesis [5]. Given the delay of response (> 1*hr*), a genomic regulation of ACTH synthesis seems a plausible explanation of the concentration profiles; although the authors propose a rate-sensitive mechanism, a threshold mechanism relative to a set point seems more plausible. Regulation seems to act mainly on the CRH reception or transduction, this being almost silenced by cortisol in basal conditions and becoming the main secretagogue when cortisol levels fall [5]. Contradictory evidence in rats suggests that GC regulation of HPA activity acts on CRH release, but not in CRH-mediated ACTH release [8]. There is also evidence of GR nuclear translocation desensitization in response to long-term constant CORT concentrations (7 days) [143]; this mechanism might underlie a rate-sensitive regulation of genomic responses at ultradian time-scales, but there’s no evidence of this. The precise mechanism of GC repression of POMC transcription is unknown [122]; GC protein-protein interactions with the CRH signaling factor NGFI-B inhibit its activitory function on POMC transcription [15, 154], but promoter deletion experiments suggest that GC acts on Tpit/Pitx1Re activation but not on NuRE activation, interference with SRC coactivators recruitment has been postulated among other possibilities [122]. However, apparently contradictory evidence suggest that GC don’t seem to inhibit ACTH synthesis and even it may increase ACTH content for high GC concentrations [142]; this apparent contradiction may come from the fact that GC interferes with CRH stimulated and ACTH release and that GC probably can not inhibit basal transcription and probably AVP stimulated transcription.

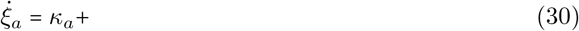
- *Regulation of receptor synthesis:* the number of V1bR receptors increase during chronic stress, while CRH receptors are densensitized and downregulated [2]. V1bR translation appears to be under tonic inhibition by upstream minicistrons and positive regulation through protein kinase C (PKC) activity [168].

**adrenal zona fasciculata/reticularis (AZFR) cells:** Contrary to other secretory cells in the HPA system, AZFR cells release steroids immediately after synthesis into the systemic circulation by diffusion [129]. Given that there’s no accumulation of the hormone in vesicles, synthesis and secretion are tightly coupled. However, as in other endocrine cells, there are two different time-scales for the biosynthesis-secretion process, one involving fast intracellular signaling and membrane depolarization acting on the scale of seconds to minutes, and the other involving the genomic machinery acting on the scale of hours [49, 48].

*Cortisol biosynthesis:* Just as corticotrophs in response to CRH, AZFR cells internal cell signaling is mainly triggered by the activation of the cAMP pathway following ACTH binding to receptors. This signal triggers cortisol biosynthesis in two phases, both slow and fast steroidogenic processes [49].

- In the first phase, ocurring within seconds to minutes, a fast ionic signaling is elicited by cAMP mediated inhibition of background bTREK-1 *K*^+^ channels, producing a cell depolarization and entry of *Ca*^++^ by activation of VGCC. ACTH inhibits outward *K*^++^ channels by two alternative cAMP-dependant signaling pathways, one involving PKA and the other Epac2, a guanine nucleotide exchange protein activated by cAMP [107]. ACTH activates cAMP production mainly through adenylyl cyclase 5/6 and adenylyl cyclase 2/4 [36]. Both cAMP and [*Ca*^++^]_*i*_ elicit fast biosynthesis by activating synthesis of steroidogenic acute regulatory (StAR) protein, stimulating cholesterol uptake and intramitochondrial transfer [49]. Contrary to corticotrophs, neuroendocrine cells and neurons, AZFR cells don’t express voltage-gated Na^+^ channels that may sustain action potentials and explain sustained and significant [*Ca*^++^]_*i*_ increase by repetitively activating VCCG channels; however, a model has been proposed where following the inhibition of background *K*^+^ channels, two opposing voltage-dependent currents are activated, one is the VCCG and the other a *K* voltage-dependent, rapidly inactivating one, such that they both may sustain action potential bursts and sustained *Ca*^++^ entry [49].
- The second slower biosynthesis processes, occurs by induction of the synthesis of the enzymes that convert cholesterol to cortisol as well as the induction of the expression of bTREK-1 channels; they are also triggered by cAMP activation and accelerated by *Ca*^++^ entry and requires hours [49, 48].

*Secretion:* By simple diffusion across the cell membrane.

Keenan et al, have proposed the following model for the transfer (dose-response) function from input signals of hormones **q** onto *i*-hormone rate synthesis:

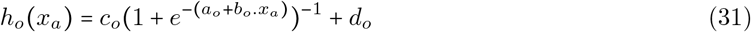

Measures of steady-state ACx total secretion in humanas are available with a resolution of some minutes. Typical parameter values can be estimated from these data have been given in [87, 88]:

| $a_o$ | $b_o$ | $c_o$ | $d_o^4$ |
| --- | --- | --- | --- |
| -4.76 | 0.14 | 11 | $0.1 \times 10^{-6}$ |

The implicit assumption is that the AZFR steady-state is reached within this temporal window, so that it is much faster than changes of concentration in blood and therefore, at the scale of 10 *min* it can be described by an instantaneous function of the delayed and transformed concentration signals.

#### 3.1.4 Inputs and perturbations

**Physical and psychological stressors:** Studies of neural by Dayas et al have suggested that stressors are processed by the brain at least in two categories, each one following a different neural path: physical stress (immunological, haemorrhage, etc) elicit neural activity in Noradrenergic cell populations of the medula in regions that are more rostral than those activated by psychological stress (restreint, noise, etc), and activates cell populations in the central nucleus of the amygdala while psychological stress activates the medial nucleus [40]; immunological stress have been also been shown to activate noradrenergic cells in the nucleus tractus solitarius (NTS) from which the information is relayed to the central nucleus of the amygdala [18]. Psychological stress information is also relayed both from the medulla directly to the PVN and indirectly from the amygdala to CRH neuron populations in the PVN [41]. While responses to immunological challenge seems to be strongly depend on the noradrenergic projections from both the VLM and the NTS [18], psychological stress seem to more strongly depend on the indirect path involving the amygdala and being less dependent on NE terminals [41]. The strength of HPA responses to stress depend on the stage of the glucocorticoid ultradian pulses during which the stress occurs, active during the rising phase and inhibited during the decrease phase (refractory period) [106].

**Acute versus chronic stress:** While ACTH concentrations increase during both acute and chronic stress, hypothalamic responses show important differences; both CRH and AVP increase in response to acute stress, whereas during chronic stress portal blood CRH concentrations and parvocellular mRNA diminish and those of AVP strongly increase [106]. In addition, during chronic stress the pulsatility strongly increases producing an apparent hyporesponsiveness to additional stress and flattening the circadian rhythm [106].

**Molecular mechanisms:** Il-1*β* applied directly to pPVN neurons facilitates pPVN cells activity apparently by interfering with afferent GABA inhibitory activity [54, 53]. NE exert different influences on two different subpopulatoins of pPVN neurons: it facilates Glu-mediated synapatic activity on some cells, via de *α*1-adreno receptor, and it directly inhibits (hyperpolarizes) other subpopulation via the *β*-adreno receptor [38]. Bacterial endotoxin LPS directly stimulates interleukin IL-6 release by mouse FS cells [109] which induces ACTH secretion by corticotropes [63]. MIF cytokine release by AP (corticotropes, FS, other) cells is stimulated by both LPS and glucocorticoids and inhibits glucocorticoid inhibition of IL-6 secretion; in corticotropes MIF secretion is also stimulated by CRH and the concentration required to release MIF is lower than that required to release ACTH [163].

**Maternal hormones:** Allopregnalonone, a neurosteroid, progesterone metabolite and modulator of GABA-A receptors, induce endogenous opiod enkephaline secretion by neurons of the NTS which inhibits NE secretion in neuron terminals in the PVN, supressing responses to stressful stimuli in rats [16]. Similarily, in women a physiological dose of estrogen can restrain cytokine and neuroendocrine responses to an inflammatory challenge [136].

**Circadian and light information:** *SCN → pPVN:* Light responding cell populations from the retino recipient ventrolateral SCN project to the pPVN (in rat and hamster) onto which they influence the HPA system rhythms [121]. Light affects these rhythms only at times where the endogenous SCN circadian rhythm is responsive (subjective night), the light induced activity of SCN neurons is coupled to phase shifts of the circadian rhythm [121]. Activation of the SCN → pPVN pathway seems to be phase dependent, with higher activity during the late portion of the subjective night [121]. However, the effect of SCN activation on the activity of PVN neurons is different for different cell subpopulations: medial and dorsal PVN neurons sending axons to the ME are mostly activated while those in the paraventricular subnucleus of the PVN projecting to the ME and those in the medial PVN sending projections to the Dorsal Vagus Complex (DVC) in the brainstem are inhibited (as well as magnocellular cells sending axons to posterior pituitary) [76]. The prescence of GABA_*B*_ receptors modulate GABA mediated inhibition [169, 108]. *Molecular mechanisms:* VIP and GIP are two of the main neurotransmitters involved in this interaction while AVP is rarely involved, but most (> 50%) contain another neurotransmitter [121]. Most of them synthetize the inhibitiory *γ*-aminobutyric acid (GABA) [121] which has been shown to act monosynptically together with the excitatory Glutamate (GLU) onto the PVN [175].

*Receptors:* Both non-NMDA and NMDA for Glu in pPVN [75]; GABA_*A*_ [75] and GABA_*B*_ [169, 108].

*Delay:* 12.6±0.6 msec (GLU), 16.6±0.6 ms (GABA) [75].

### 3.2 Implementation

The hybrid automata implementation with SpaceEx imposed some constraints on the kind of models that can be represented and executed, the most important being that only affine dynamics are supported. However, biological processes show plenty of nonlinear mechanisms that underlie their special dynamic properties, such as multistability [159]. In the ODE system of equations describing the dynamics of the elementary functional units, we find mainly three kind of nonlinearities in addition to the affine ones: i) steep sigmoideal functions and ii) products and iii) powers. In order to implement them in SpaceEx, we must find piece-wise linear or affine approximations. We must however be careful and take into account for the analyses of the results, that different approximations to nonlinear ODEs may show very different behaviors that not always correspond to the behavior of the original nonlinear ODE system [134].

### 3.3 Piecewise linear or affine approximations

**Hill function:** as a first approach, we approximate these functions by step functions, where the threshold of the step function is given by the abscissa corresponding to the maximum slope, such that the flow (evolution) becomes affine or constant in each location:

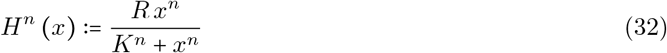

**Approximation with step function:**

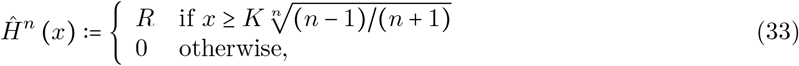

The model has X continuos state variables (free CRH, ACTH and cortisol blood HC, accumulated CRH, ACTH in synaptic vessels, non free protein-bounded cortisol, CRH neurons FR) and 4 additional variables controlling hormone accumulation and release in the CRH neurons and AP. It is described by the following equations (For *i* ∈ {*c*, *a*, *o*}, the letters designate both the signal (hormone) and the structure originating the signal (*c*=CRH (ME), *a*= ACTH (AP), *o*= cortisol (ACx))):

### 3.4 Corticotroph fast secretion unit

**Elementary units:**

- Simple receptor: deppending on kinetic constants, the basic hormone receptor responds either instaneously (if rise and fall times are much faster than the time scale unit), or according to:

## 4 Discussion

**Concentration data:** A first consideration that is rarely taken into account in most models is about the data acquisition. In fact, the data made available for us by F. Roelfsema is sampled every 10 minutes. However, more detailed measures of CRH and ACTH taken in horses every 20 to 30 sec have shown the existence of oscillations with with mean interpeak intervals of approximately 5 min. Signal processing considerations suggest the signal to be filtered at half the sampling frequency in order to supress aliasing (high frequency power being mapped to low frequencies, distorting the data). Once the signal is measured it becomes very difficult to distinguish real low frequency oscillations from high frequency ones aliased into the low frequency range. Therefore, we must assume that part of the fluctuations seen in the data are indeed due to aliasing from higher frequencies.

**Ultradian Oscillations** An important question that arises from the analysis of HPA system models, is which is the origin and importance of the ultradian pulsatory patterns. In fact, in Keenan’s model the pulsatory activity is modeled by a stochastic process that generates a series of dirac pulses which are then convoluted with an hypothetical pulse waveform. Contrary to other models, Keenan et al.’s model is not closed, the parameters of pulse generation process are free and therefore the caracteristics of the ultradian oscillations do not result from the endogenous dynamics of the HPA system but from the statistics of the pulses generated by the renewal process and from the parameters of the hypothetical pulse waveform with wich it is convoluted [87]. This modeling choice is consequent to the fact that the model is used to estimate the pulse parameters and not to study the system dynamics. However, as nothing is said about how this process is controlled and the parameters of this process are effectively disconnected from the rest of the network, the pulses are supposed to be produced exogenously to the network dynamics. Nevertheless, alternative modeling studies propose that ultradian oscillations are endogenously produced by the dynamics of the network. In particular, the work of Lendbury and Pornswad, suggest that the pulsatory activity is indeed produced by the delayed negative cortisol feedback on ACTH synthesis and the one from ACTH concentration on CRH synthesis [102]. Recently, an empirical study appeared in the litterature that supports, indeed, the structure of the second model [141].

**Rate-sensitive feedbacks:** several works have shown the existence of a fast rate-sensitive feedback both for CORT secretion after stress stimulus (CRH release) [106] and for basal CORT secretion (without stress stimulation) [10], but the precise mechanism underlying this feedback is unknown. Keenan et al have proposed a model taking into account this fast feedbacks, but they didn’t propose any model for the mechanism. One possibility is that the sensitivity to cortisol concentration rate involves mainly the different affinities of GR and MR to GCs [106]; however, it is not at all clear how this may happen. A possible mechanism, that has not been evoked, is the one composed of i) a “high pass” filtering exerted by CORT binding to albumin other blood proteins, together with ii) a differential activity of intracellular and membrane-bound receptors (which has indeed been evoked in [10]). The hypothesis being that in normal “basal” conditions, there’s only a small, slowly changing, amount of available free cortisol that is rapidly and strongly sequestred by the high concentration and affinity of intracellular MR and GR receptors, which would not participate to the fast response influence slow transcription processes; in this case, there will be no enough free cortisol to stimulate membrane-bound receptors. On the contrary, during acute secretion or rapid injections, the amount of free cortisol augment considerably for a short time, rapidly saturating available intracellular receptors and activating the membrane-bound receptors involved in the fast feedback.

## A Appendix

## A.1 Basic mechanisms

## A.1.1 Signal reception and transduction

Several types of stimulus are functional in neuro-endocrine systems. Peptide secretagogues such as CRH and ACTH act as ligands, binding either to membrane G-protein Coupled Receptors, which in turn activate their attached G proteins (breaking them in two active, moving free, subunites: G_*α*_ and G_*β*_/γ), or to ion channels coupled receptors, among other mechanisms. Steroids such as CORT either diffuse into the cell where they bind and activate intracellular receptors, which enter the nucleus and bind DNA promoters, or bind membrane-bound receptors or directly interact with the membrane modifying its physicochemical properties [50], glucocorticoids bind both to glucocorticods receptors (GR), with low affinity (dissociation *t*_1/2_ ~ 5 min), and to mineralcorticoid receptors (MR), with high affinity (*t*_1/2_ ~ 45 min), but may also use other transduction pathways [50]. Paracrine signals and electrical signals act by direct contact between cells [4]. Electrical signaling is not only obseved in neurons and neuroendocrine cells, but also in endocrine cells such as those in the anterior pituitary where action potentials and gap junctions have been observed [156].

**Table 1:**
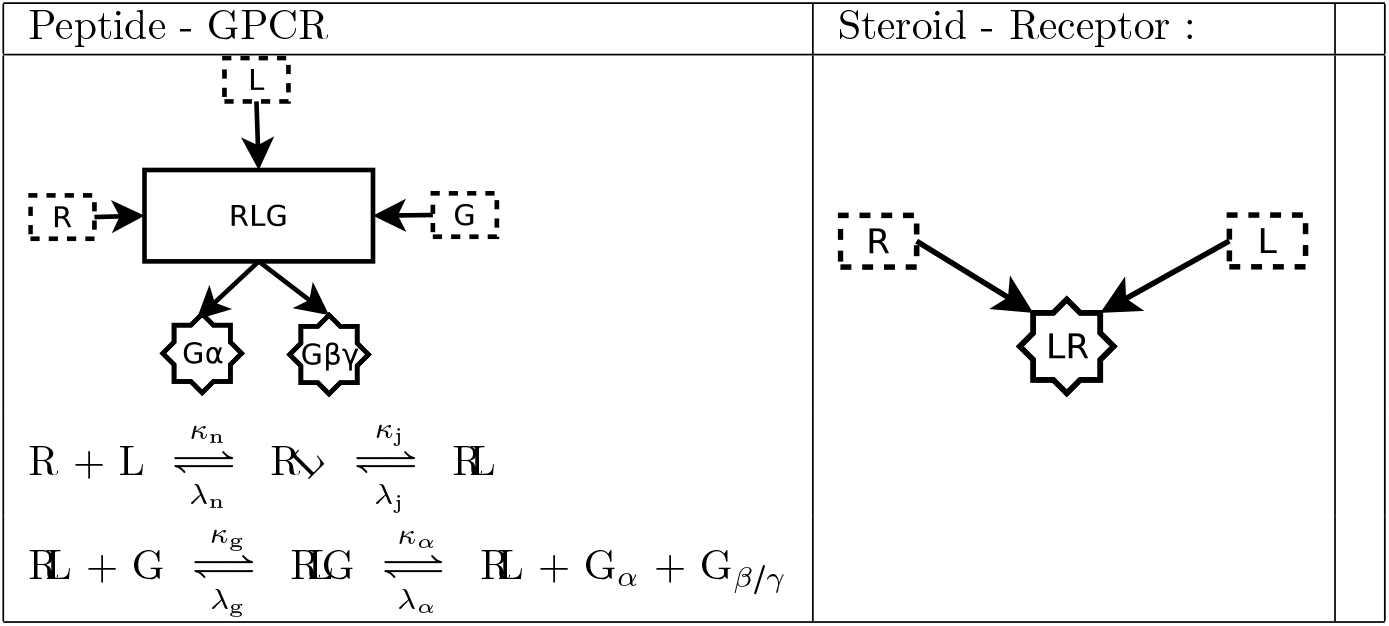
Peptide - GPCR mechanism:

- *Peptide - GPCR mechanism:* following [78] we assume the a two-domain model for the ligand - receptor reaction (see Table 1), first the ligand C-terminal binds to the receptor N-domain 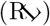, increasing the effective affinity of the ligand N-terminal for the receptor J-domain; then the ligand N-terminal binds the receptor J-domain forming the 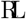 complex. If the receptor is in a G-protein bound state then ligand binding leads to the liberation of the protein subunits (G_*α*_ and G_*β*_/γ); G_*α*_ is then free to bind to GTP becoming active. Denoting *r* =[R], *l* =[L], 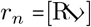, 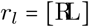 the dynamics are given by:

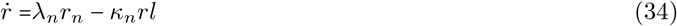

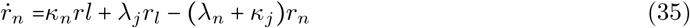

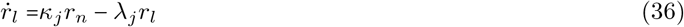

assuming a constant total concentration of GPCR (*r*_*tot*_ = *r*+*r*_*n*_+*r*_*l*_) one finds the following equlibrium relationship for the receptor activated by the ligand *l* (*r*_*l*_):

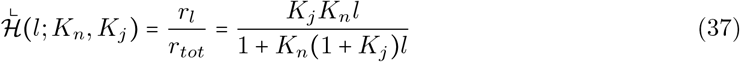

where *K* = *κ./λ* are the equilibrium association constants. It has been shown that when the receptor is in a G-uncoupled state it shows very low affinity for the ligand, whereas when it is in the coupled state its affinity is strongly increased, and for very strong affinities the response may be simplified to the one of a simpler receptor (as in (40)). *Desensitization:* It has been shown that GPCRs binding by its ligand triggers phosphorylation of the receptor inducing desensitization and internalization [132].
- *Steroid - [GR,MR] mechanism:* MR binding activation: [139][81]. Glucocorticoids bind to MR and GR, and mineralocorticoids to MR, and both may also bind directly to the membrane and change its properties, triggering both genomic [125] and fast nongenomic actions (within 15 minutes) [50]. Signal transduction may use different mechanisms [50], here we only take into account the MR and GR activation. Both receptors may be located either intracellularly or associated to the membrane and activate different signal pathways [65]; we assume no differences in the mechanism for the first signal transduction step of receptor activation that is given by:

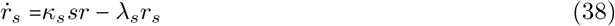

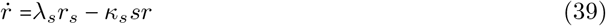

which gives at equilibrium:

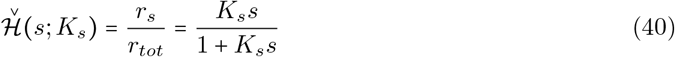
for *K*_*s*_ = *κ*_*s*_/*λ*_*s*_, *r*_*s*_ the activated receptor, and *r*_*tot*_ = *r* +*r*_*s*_ constant. It has been shown (in hippocampus cells) that GR availability is large at the cytoplasme [106] so we may assume that GR-mediated responses to glucocorticoids depend only on the incomming signal and no on receptor availability. Due to their different affinities to glucocorticoids, it is suggested that GR and MR mediate different phyisiological functions involving glucocorticoids, while GR may be involved in fast recovery of HPA responses to acute stress, MR may be involved in homeostasis [106].

## A.1.2 Intracellular signaling

two pathways involing proteins kinases, PKA and PKC, have been shown to modulate exocytosis in almost all regulated secretory cell [120]; these kinases have been shown to regulate multiple aspects of secretion by phosphorylation of multiple protein targets [110, 120]. However, other PKC- and PKA-independent pathways has also been shown to mediate exocytosis regulation [147], one of them involves an intracellular [*Ca*^++^] signal.

- *Gs*_*α*_ - *adenylate cyclase (AC) - cAMP - PKA:* this signaling pathway consist of four basic reactions: i) activated (GTP-bound) *Gs*_*α*_ proteins subunits bind and activate the enzyme adenylate cyclase (AC); ii) activated AC catalyzes conversion of ATP to cAMP; iii) cAMP bind and activates the enzyme protein kinase A (PKA) catalytic subunits, which activate different sets of target proteins including membrane ionic channels and transcription factors by phosphorylation; iv) PKA phosphorylates phosphodiesterases (PDE) [4]. Several AC isoforms are expressed in mammals, all of them activated by *Gs*_*α*_ but they are also activated and/or regulated by other molecules that differ across isoforms, resulting in complex and rich possibilities of regulation by different signals in different cell types [68]. No precise available information has been found about the particular isoforms expressed in neuroendocrine or pituitary cells; however, it is known that AC5/6 and AC9 are expressed in the adrenal gland [42, 36, 68] and that AC8 is expressed in hypothalamic nuclei and therefore is probably related to neuroendocrine functions [42]. It has also been suggested that the signal amplification from receptor to AC is neglibible due to the low number of AC molecules, which represents a limiting factor [6, 135]. Both cAMP synthesis and cAMP break down by phosphodiesterases are very fast making this process very rapidly responsive (seconds) [4]. Complete models of this pathway for yeast in response to glucose were recently published [173, 22], which suggest that the most important steps in the pathway are ii) and iii): whereas the first reaction is fast and its gain is almost unity ([22]). Following [173], starting from a Michaelis-Menten formulation for cAMP synthesis and for its degradation by PKA phosphorylated PDEs, and assuming some simplifying hypotheses, we may describe cAMP dynamics as follows:

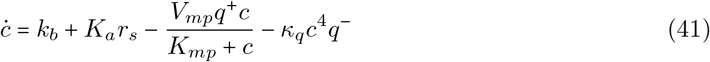

where *c* = [*cAMP*], *q*^+^ and *q*^−^ are the activated (free cataylytic unit) and inactive (complex) PKA, respectively; the first term, *k*_*b*_, represents a basal AC activity; the second term represents cAMP synthesis, which closely follows CRH receptor activation; the third term represents cAMP hydrolysis by PKA phosphorylated PDE, *V*_*mp*_, *K*_*mp*_ are the effective Michaelis-Menten constants for the PKA-PDE enzymatic chain; the third term corresponds to the amount of cAMP that binds the PKA inactive complex. ATP is assumed to always available in greater concentrations than cAMP, such that the maximum rate of cAMP production is always reached; in addition, this maximum rate depends on the activated AC which itself is assumed to reflect (with a unity gain [135]) the number of activated receptors. PDE phosphorylation by PKA is assumed to be fast and PDE is assumed to be always available, and the receptor to AC activation chain to be fast and with unity gain. However, the parameters for this reactions are unknown.

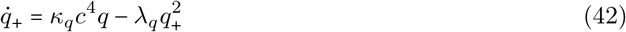 The cAMP accumulation at steady-state follows the experimental logistic function whose parameters are given in [78] (*EC*_50_=0.042 *nM*). The amplification factor is unkown (to our knowledge) for this particular signaling chain; however, it is known that the gain can be very high (tenfold or more for serotonin responses [4] (10^−6^ *M*/5×10^−8^ *M*)); this information may be used to estimate the parameters of this module ^5^.
- *G*_*q*_*α phospholypase C (PLC) → IP3 / DAG → PKC:* Other kind of G proteins respond by activating the enzyme phospholypase C (PLC) [**?**]. There are many classes of PLC isozymes, the PLC-*β* type are activated by *G*_*q*_ proteins; the more widely expressed [**?**] and in particular in AtT20 cells [**?**] (which are usually used to model corticotrophs) are PCL-*β*1 and PCL-*β*3 which bind *G*_*q*_*α* with a dissociation constant of (*Kd*) of 40–60*nM* [**?**]. The activated PLC enzyme synthetizes IP3 and DAG from plasma membrane phospholypids; DAG then activates protein kinase C (PKC) and activated PKC further propagates the signal by phophorylation [4]; IP3 induces the release of intracellular stored *Ca*^++^ by activation of IP3 channel receptors (IP3R); the increase on [*Ca*^++^]_*i*_ triggers multiple intracellular processes after binding to target proteins [166, 4].
- *GR and MR nuclear translocation:* activated receptors dissociate from a regulating multiprotein complex and enter the nucleus where they can directly bind DNA as an homodimer and activate or repress transcription, or act by other indirect mechanisms [125].

## A.1.3 Electrophysiological mechanisms

- *Spontaneous electrical activites* are indirectly modulated by chemical stimulus by the activation of intracellular signaling pathways. In this way, a single molecule may act both as a direct secretagogue and as a modulator of spontaneous electrical activities leading to secretion [156], in addition to a possible action on the synthesis of the secretory products.
- *I*_*h*_ *currents:* hyperpolarization-activated and cyclic nucleotide-gated (HCN) channels have shown to be active both in hypothalamic pPVN cells [137] and in AP endocrine cells [156]. In addition, CRH has been shown to depolarize both kind of cells [92, 93, 137] and to enchance I_*h*_ currents in pPVN cells [137]. This rises the possibility of local oscillatory electrical activities in both kind of cells dependent on I_*h*_ currents. However, HCN seem to have a negligible role in AP pacemaking electrical acitivity [156].
- *K++ inward rectifying channels:*
- *Voltage Gated Calcium Channels (VGCC):*

## A.1.4 Calcium ER storage and release

there are two kind of channels involved in regulation of intracellular stored *Ca*^2+^ release, Inositol trisphosphate receptor (IP3R) and ryanodine receptors (RyR), a main difference between them is the presence of the IP3 binding site in IP3R [56]. IP3 activates Ca^2+^ release by binding to its receptor, which is a *Ca*^2+^ channel, and activating it by reducing the inhibitory action of high [*Ca*^2+^]_*i*_ [113]. The IP3R shows (*Kd* ~ [10 − 100] *nM*) and a halfmaximal flux activatation has been observed at [*IP*3] ~ [40 − 80] *nM* [56]. In addition to activation by IP3, the IP3R has binding sites for allosteric regulation by other intracellular signals or states, including [*Ca*^2+^]_*i*_, PKA and PKC, among others [56]. Differential biophysical properties among IP3R types, in particular difference on [*Ca*^2+^]_*i*_ regulation, underlie different [*Ca*^2+^]_*i*_ signal waveforms triggered by its activation [56, 176] and therefore the different AVP responses observed in corticotrophs [34] may result from differences on IP3R types expression. The total amount of *Ca*^2+^ in the lumen may be > 1 *mM*; the concentration of free Ca2+ has been estimated to be between 100 and 700 *M*, whereas, the concentration in the cytoplasm of unstimulated cells is between 50 and 100 *nM*, [56].

The inverse process: *Ca*^2+^ storage in the Endoplasmic Reticulum (ER) is regulated by a complex mechanism involving STIM and Orai proteins, the former act as sensors of ER stored *Ca*^2+^ depletion and the latter ones form a *Ca*^2+^ selective channel [60].

## IP3R channels

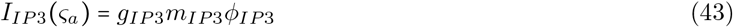

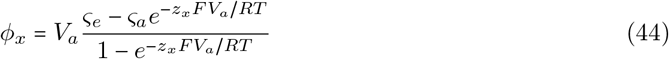

## A.1.5 Secretion

- *Regulated vesicle secretion:* (see figure 10) in neurons and neuroendocrine cells, as in most endocrine other secretory cells, is a process triggered by an increase on intracellular Ca^++^ following a chemical or electrical stimulus [46]. In endocrine and neuroendocrine cells it is known to be a slow process with long latencies, of the order of seconds or minutes, compared with neurotransmitter release, which occurs in a period of the order of msecs [146, 46]. A common pattern shared by different kind of cells is the storage of the secretory product in intracellular granules or vesicles [46], but some secretions occur without vesicle formation by simple diffusion of the products across the cell membrane. Differences among cells in Ca^++^ - secretion coupling depends on the density of Ca^++^ channels, the distance between the Ca^++^ channels domain and the release-ready secretory vesicles or granules, which is much bigger in pituitary cells than in synapses [155]. Ca^++^ occurs either by Ca^++^ channel opening and inflow or by intracellular Ca^++^ release and can be elicited and modulated by multiple mechanisms [172, 155]. The stimuli-induced and Ca^++^ dependant fusion of granules to the special sites in the cell membrane called *porosomes* across which the product is expulsed to the exterior of the cell [83]. Secretory vesicles are stored in different pools, some vesicles being near calcium channels and ready to be released and others used as a reserve [131]. Often, in endocrine cells, there is only a partial discharge of secretory products in response to a stimulus (kiss-and-run phenomena) [83, 172]; consequently, the peak amplitude of the [Ca^++^]_*i*_ increases progresively as the length of the stimulus increases [165, 155]. Several seconds (~5sec) of depolarizing stimuli are thus needed to completely deplet the ready-to-be-released pool, after which the pool slowly recovers to reach the maximum pre-stimulus level [165]. However, the size of these pools and probably the velocity of replenishement may be modulated by several factors, such as the available intracellular concentration of hormone precursors, and some may also underlie a mechanims of plasticity (learning) phenomena [172]. In neurons, due to the small distance from vesicles to Ca^++^ domains, a single action potential and even a single calcium channel opening can elicit neurotransmitter release [153, 170, 156]. The proposed Ca^++^-sensor synaptotagmin (Syt) seems to prevent vesicle-membrane fusion and Ca^++^ seems to ease this process by binding to Syt [172].

**Figure 10:**
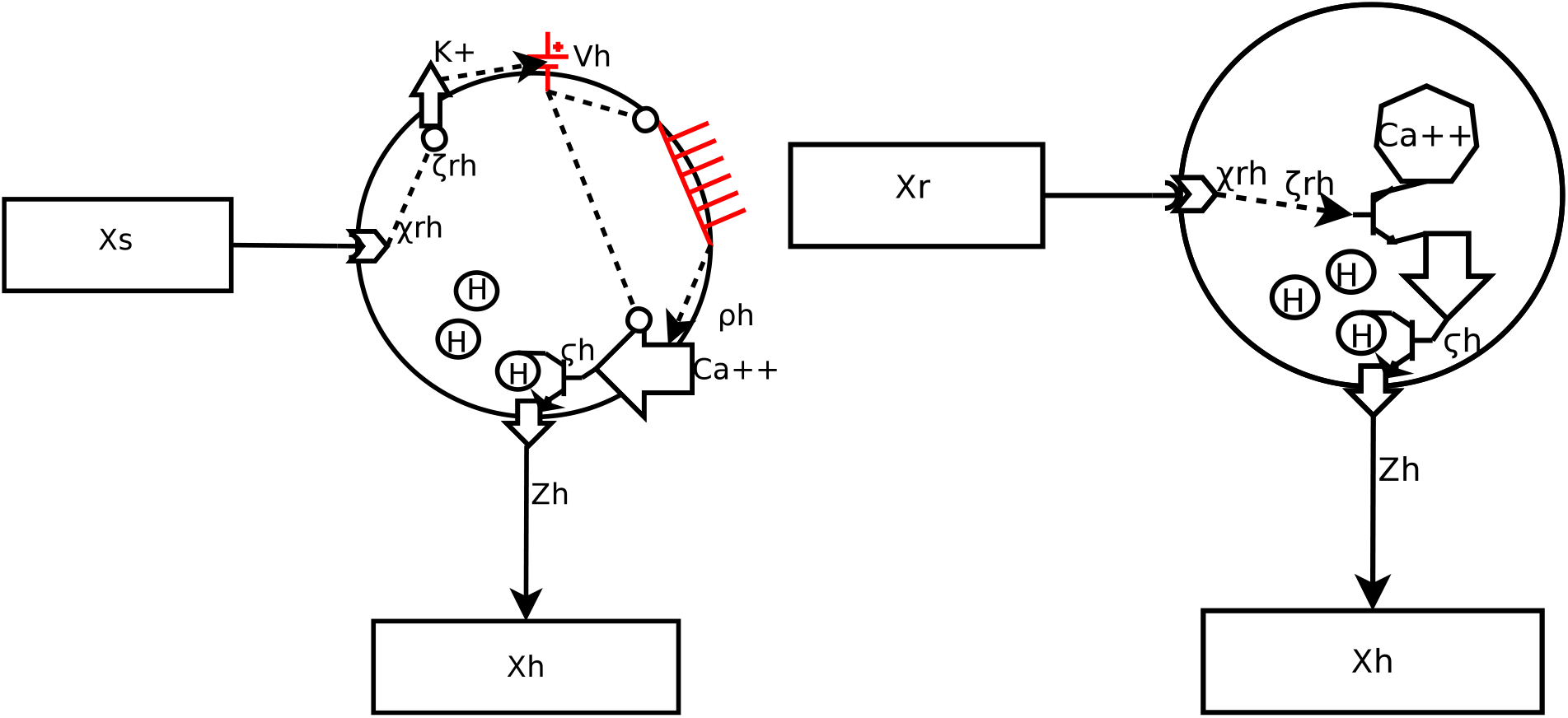
Secretion control may occur in at least two (non-exclusive) alternative pathways. In the first (left), receptor activation by the releasing factor (*x*_*r*_) activates an intracellular signaling pathway (*ζ*_*rh*_) that modifies some ion current (e.g. K^+^), modifying the membrane potential (*V*_*h*_), which facilitates action potential and increase firing rate (*ρ*_*h*_), depolarizations controls a VCCG allowing *Ca*^++^ inflow and *ς*_*h*_ = [*Ca*^++^]_*i*_ trigger vesicle fusion and hormone exocytosis, increasing secretion rate(*z*_*h*_). In the second (right), intracelluar signaling triggers internal *Ca*^++^ release, with the same results but releasing internal *Ca*^++^ stores, without the need of VCCG channel activation. In [144, 145] the following kinetic model was proposed to describe Ca^++^-regulated vesicle exocytosis:

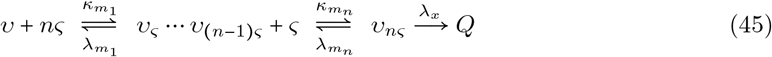 Where *ν* is the vesicle concentration, *ς* is [*Ca*^++^]_*i*_ and *Q* is the concentration of released quantums of hormone (*Q* = [*q*], *q* being the mean hormone amount released by a single vesicle during a single release event). However, in these equations the quantity of available vesicles is assumed constant, which is not relistic, so that they are only valid in the very short term; therefore, we modified it to take into account readily-available vesicle pool depletion.

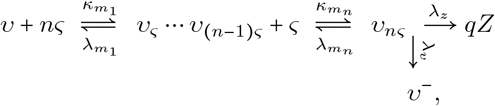

where *z* is the concentration of secreted hormone, *q* is the number of molecules secreted in each release event, *ν*^−^ is the concentration of depleted vesicles, *ν*^+^ = *ν*_*nς*_ is the concentration of fully activated vesicles, and *ν* is the concentration of readily-available vesicles. The dynamics are then given by:

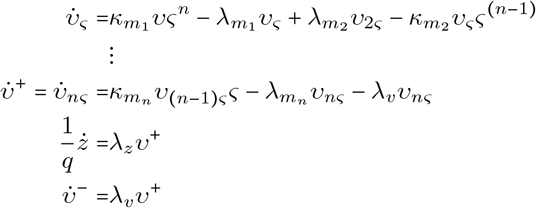

and the total number of vesicles 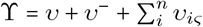 is assumed to be constant (i.e. a parameter of the system, both *θ* and ϒ depend on the output of the slower hormone synthesis and internal vesicle transport system, we don’t take into account the pool of full but not readily-available vesicles). The rate of vesicle depletion is assumed to be determined only by the amount of hormone that has been secreted, such that:

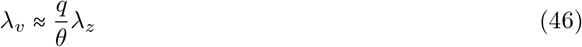

where *θ* is the approximate quantal availability of each vesicle before any release, which is a parameter of the system. Additionaly, we may assume a high cooperativity of [*Ca*^++^]_*i*_ in inducing vesicle contents release, as it has been shown to be the case for synaptic vesicles [145] and is compatible with hill equation curves fitted to corticotroph data [165], so that we may assume cooperative Hill dynamics for secretion:

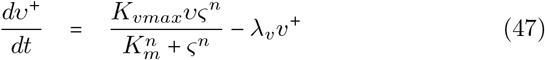

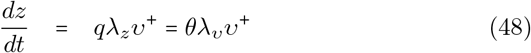

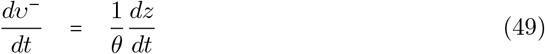

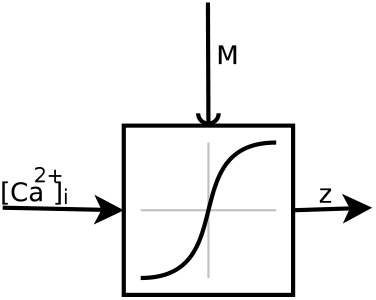

## A.1.6 Vesicle recyclement and readily-available pool replenishement

Granule biosynthesis and transport to secretory sitesis is an extremely complex process [154, 130, 131]; prohormones/proneuropeptides, among other factors, are able to trigger the process [130]. It has been suggested that endocytosis has an essential role for this process in the recycling of vesicles needed for continuous secretion [47]. However, at least in corticotrophs, recycling of vesicle contents seem to be negligible [127] so we may neglect partial secretion and recyclement of contents. Endocytosis and exocytosis are tightly coupled in corticotrophs [101, 127] and inhibition of endocytosis in chromaffin cells results in damped secretion [47]. In addition, exocytosis induces the transcription of proteins needed for granule sythesis and there is always an excess of granules being transported to the secretory site, which are either recycled or captured to rapidly replenish the readily-available pool [130]. Therefore, we assume that the process of refillement of the readily-available vesicle pool reuse of endocytosed vesicle membranes and that an homeostatic equilibrium exists between new vesicles created and available, new reabsorbed, old reused and old degradated, such that we may neglect vesicle creation at basal conditions.

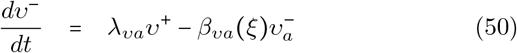

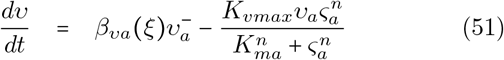

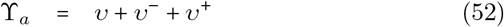

where *β*_*νa*_ represents the rate at which new filled vesicles are created, which is a function of is the prohormone/proneuropeptide concentration *ξ*. Assuming the process to happen in a cooperative fashion with respect to *ξ* and neglecting transport to secretion sites (probably important), it may be given by:

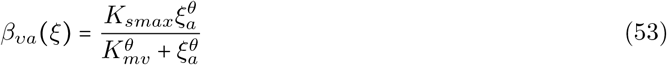

where *η* = *θ* × *N*_*A*_ is the mean number of molecules of prohormone in each vesicle (*N*_*A*_ being the Avogadro number). However, as *η* is typically a big number (see A.2.5) this hill function is very steep, so under an important basal synthesis *β*_*νa*_(*ξ*) would basically reach constantly *K*_*smax*_ in normal conditions.

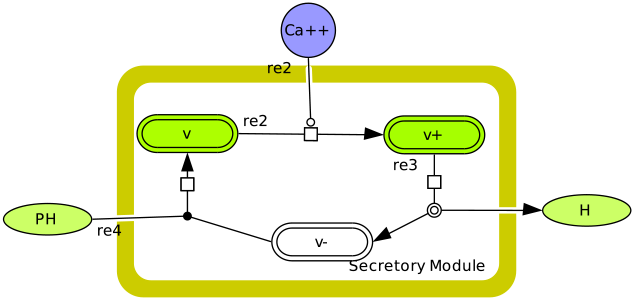

## A.1.7 Synthesis

(see figure 11)

**Figure 11:**
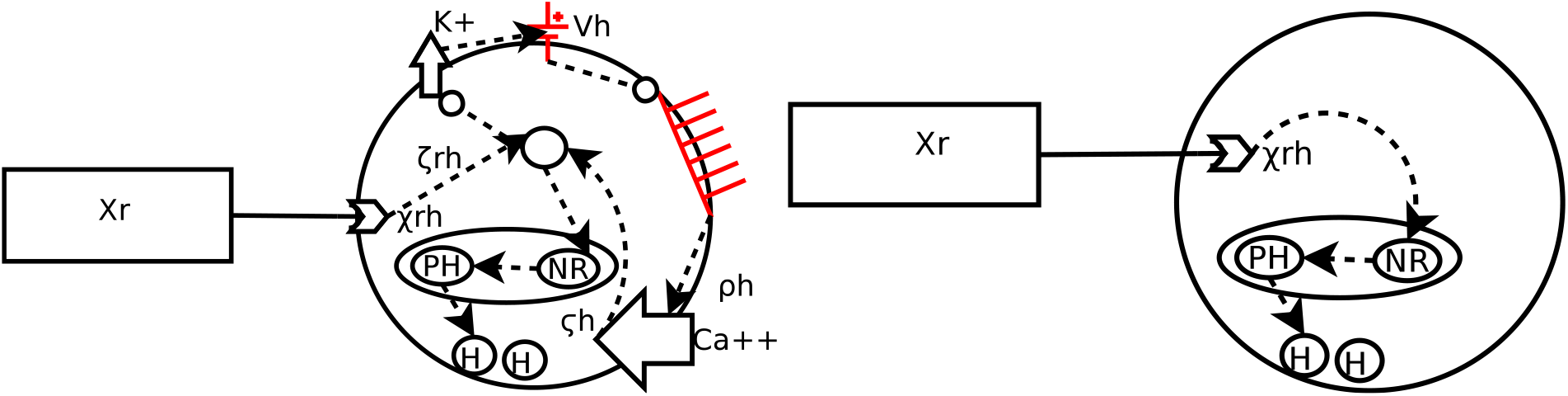
Synthesis control occurs also in two alternative pathways. In the first (left), membrane receptor activation by the releasing factor (*x*_r_) activates an intracellular signaling pathway (*ζ*_*rh*_) that acts in two different processes, either it enter the nucleus where it activates/regulates hormone transcription, or it takes the same path shown before for secretion (see figure 10) allowing *Ca*^++^ inflow and *ς*_*h*_ = [*Ca*^++^]_*i*_ activates the hormone transcription process. In the second (right), intracellular receptors are activated and translocated to the nucleus, where they regulate transcription.

## A.1.8 Blood hormone transport

Blood transport may be represented by a simply linear advection-diffusion-decay equation in a single spatial dimension (along vessels length) with an additional term representing the degradation of the hormone during the transport and that we assume proportional to the concentrationo of the hormone at any point in the circulation vessels:

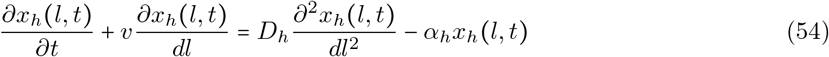

Where *ν* is the velocity of the blood stream, *x*_*h*_ are the concentration of hormone *h*, and {*D*_*h*_, *α*_*h*_} are its diffusion and degradation constants in blood, respectively.

The solution for this equation for an impulse of concentration 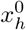 at (*t, l*) = (0, 0) is given by:

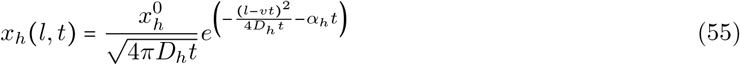

However, we are not interested in the solution along the blood vessel, but only at sinks sites for each hormone (with respect to the their source sites). In addition, given that the distances between the endocrine glands is much bigger than the size of the glands themselves we may simplify the solution assuming that the dispersion 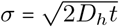 is almost constant along the gland surface receiving hormone *h*. The signal available for a sink cell is therefore given by:

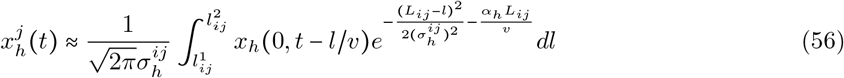

where 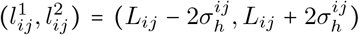, 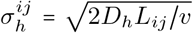 and *L*_*ij*_ is the distance between the source (*i*) and sink (*j*) cells. The integration is limited to the maximum dispersion window of length 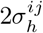, it is assumed that at any single point in the receiving gland a cell measures the dispersed and delayed instantaneous concentrations of hormone secreted from the source passing through a cross-section of the vessel.

We neglect the complexity of the blood circulation network and more complex phenomena due to the varying size of the vessels, diffusion and advection in other directions, compliance of the vessels, recirculation, etc. However, this are important parameters ([126, 70]) that must be taken into account in future work.

## A.1.9 Hormone accumulation during propagation

during transport, hormones bind to proteins such only a fraction of hormone is free to act on targets immediatly. Denoting the fractions of protein binded-hormone for *N* binding proteins as {*x*_*hn*_}_*nϵN*_, the transport equation, and assuming a much bigger concentration of binding proteins than that of the hormone, the transport equation is modified as follows:

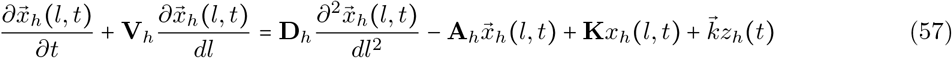

Where (e.g, for two binding proteins):

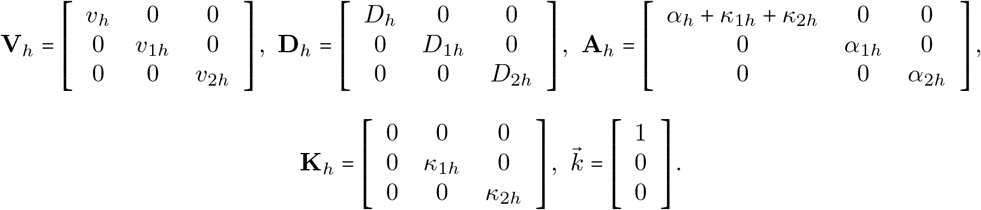

and *z*_*h*_(*t*) is the rate of hormone production at the source, {*κ*_*nh*_, *α*_*nh*_} are the rates of binding to and dissociation from the *n* protein, respectively.

## A.2 Parameter values/estimations

## A.2.1 Basic parameters

| Quantity | Value |
| --- | --- |
| mol | $6.02214078 \times 10^{23}$ elements |

## A.2.2 Molecular properties

| Molecule | Molar Mass (g/mol) | references |
| --- | --- | --- |
| CRH | 4759 | (http://www.uniprot.org/uniprot/P06850) |

## A.2.3 Receptor and transduction models

## G-RF1 response to CRH

There are several sources of information concerning the dynamics of ligand binding to the receptor [93, 174, 14, 78] and of the whole receptor system including G-protein activation [14], from where parameter estimates for the whole receptor system can be drawn. CRH binds G-CRF1 with high affnity [174, 27, 78], reaching steady-state in ≈ 30 *min* at 22°C, and eliciting fast electrical responses (onset time < 1 *min* [93]); rates are temperature dependent, incresing as temperatures increases. However, in the (G-protein) uncoupled state, both affinities of the N and L terminals (see A.1) are very weak (*K*_*n*_ ≈ 1/(56*nM*) and *K*_*nj*_ ≈ 0.1 (*K*_*c*_ ≈ 1/(51*nM*))) while for the G-coupled receptor (G-CRF1) they strongly increase (*K*_c_ ≈ 1/(0.21*nM*),*K*_*cj*_ ≈ 266) (values for rat and human corticotrophes are similar) [78] (*K*_*c*_ = *K*_*cn*_(1 + *K*_*cj*_) is the macroaffinity constant). Given that the electrical response to CRH (in rat corticotrophes) is long-lasting after stimulus offset (> 10 *min* after CRH washout) [93], one may estimate *λ*_*c*_ or at least its order of magnitude, and from it estimate *κ*_*c*_:

- *λ*_*c*_: starting at equilibrium, at the end of a long saturating step CRH stimulation (as done in [93, 92]) the dynamics of activated CRF1 are given by 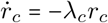, thus 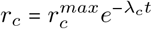. Assuming a small threshold of response of the cAMP-PKA signaling to the G-CRF1 signal at 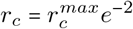 and a long-lasting response (> 10 *min*) [93, 92]: 2/*λ*_*c*_ ≈ 10 *min* ⇒ *λ*_*c*_ ≈ 0.2/*min* (it has been shown that the intracellular cAMP-PKA dependent signaling of the CRF1 response is very sensitive, responding maximaly to very small cAMP concentrations (ref)).
- *κ*_*c*_: *κ*_*c*_ = *K*_*c*_λ_*c*_ ≈ 0.95(*nM min*)^−1^ (using G-coupled CRF1 *K*_*c*_ = 1/(0.2 *nM*) [78] and *λ*_*c*_ = 0.2/*min*)
- *τ*_*c*_ (*rise characteristic time*): Assuming a step saturing CRH stimulus of magnitude *c*_*s*_, the dynamics are given by:

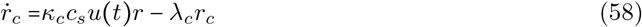

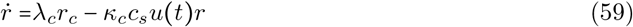 Applying the Laplace transform and assuming a constant total number of receptors *r*_*max*_ the dynamics are given by:

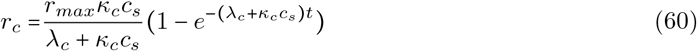

thus *τ*_*c*_ = 1/(*λ*_*c*_ + *κ*_*c*_*c*_*s*_) ≈ 0.05 *min* or 0.1 *min* for *cs* 10 *nM* and 20 *nM*,respectively.

**Table 2:** Estimated parameter values for reception and transduction of CRH, AVP and GC in corticotrophs.

| Parameter | Value | Source | Comment |
| --- | --- | --- | --- |
| $\kappa_c$ | $0.95 (nM, min)^{-1}$ | A.2.3 | Unreliable estimation, lack of data |
| $\lambda_c$ | $0.2 min^{-1}$ | A.2.3 | Unreliable estimation, lack of data |
| $\kappa_v$ | $2.1 \times 10^{-2} (nM, min)^{-1}$ | [61] | Experimental estimation (rat pituitary) |
| $\lambda_v$ | $0.17 min^{-1}$ | [61] | Experimental estimation (rat pituitary) |

## V1bR response to AVP

Binding and dissociation constants for rat pituitary are given in [61].

## GR and MR response to GC

- Binding affinities: Ligand affinity of GC for GR much lower than for MR 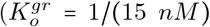, 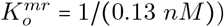[140]. However, no other data was found available in order to estimate *κ*_0_ and *λ*_o_ for GR or MR from this affinity values.
- : Activation: Cortisol binds to the human MR (hMR) with the same affinity as aldosterone, but is less efficient than aldosterone in stimulating the hMR transactivation [71]. Cortisol transactivation of MR: *EC*_50_ = 10 *nM*. Aldosterone transactivation of the MR LBD (ligand binding domain): *EC*_50_ = 1 *nM*, and for GR LBD: *EC*_50_ = 0.3 *μM* [139].

## cAMP-PKA

From the model in [173] 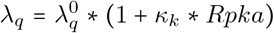 *λ*_*q*_ = 19.9 * (1 + 2.2 * 10^4^ * 1 * 10^−4^), this model includes another molecule (Krh) that is known to represses PKA activation, we take as constant the concentration of this molecule taking its maximum value to be 1 × 10(− 4)*mM* as is seen from simulations [173]. *Rpka* = 1 * 10 (− 4)

## *G*_*q*_α →PLC→IP3

*G*_*q*_*α* bind PCL-*β*1 and PCL-*β*3 with a dissociation constant of (*Kd*) of 40–60*nM* [?]

## A.2.4 Ionic Channels

## IP3R channels

## A.2.5 Regulated exocytosis mechanisms

**Porosome + Readily-Releasable Vesicle Pool:** (summary in table 3)

- *Vesicle number (ν_a_):* on one hand, electrical measures of exocytosis in rat corticotrophes *in vitro* [165] show that the membrane capacitance contributed by 1 vesicle during exocytosis is ≈ 1.3 *fF* (diameter ~ 200 *nm*), and the maximum increase on cell surface under a stimulation that deplet all be: *M* ≈ 6.12 *pF* ≈ 4708 *ves*. On the other hand, vesicle density measures on thin AtT-20 (mouse tumoral corticotroph) cells slices (50*nm*) (restricted to processes tips, where most vesicles are found) show a juxta-membrane vesicle density in of 1.5 ~ 2 *ves*/*μm*, and a total of 7.5 ~ 10 *ves*/*μm*^2^ (per tip) [171], such that for typical cell diameters ([400, 1020]*μm*, with a mean of ~500*μm* [165]), and assuming a similar density of vesicles for AtT-20, rat and human cells, some thousands of available vesicles per cell is a plausible quantity. No data specific for human cells is available, we may assume similar numbers as those in rats: *ν* ≈ 5000.
- *Exocytosis rate:* in [165] the maximum rate was found to be ≈ 2140 ± 325*f F/s*; assuming total release of contents for each event and a vesicle capacitance of 1.3 *f F*, the maximum rate of vesicle contents release is *K*_*vmax*_ ≈ 1646 ± 250 *ves/sec*. However, it has also been shown that the readily-releasable vesicle pool is depleted
- *Quantal size and vesicle content:* different measures of vesicle content are available in the litterature, according to Sobota et al. [151](citing [128] and others), granules in (rat) corticotrophs contain between 3000 and 10000 copies of POMC product, concentrations of ACTH and other hormones in granules range between 10 to 42 mM, and protein content concentration between 100 and 150 mg/ml. Mean granule diameters measures range between 90 *nm* [35] to 200 *nm* [165] (citing [28]) in rat and 356.8 ± 30.5 in human ([64]). However, this numbers are not coherent among them and thus we may suppose that some measures might be not very accurate or that the differences are related to different experimental conditions; that is: assuming the lowest ACTH concentration (10 *mM*) and a granule of 200 *nm* diameter, the number of molecules is: *V ol*_*granule*_×10×10/*mol m*^3^×*N*_*A*_ ≈ 2.5×10^5^, which is 20 times higher than the highest number of copies of POMC product reported for rat pituitary cells [151](citing [128]). Thus for our purposes we may assume for a human corticotroph vesicle the smaller reported concentration (10 *mM*), which for a granule diameter of 350 *nm* gives: 2.2449×10^−12^ *μmol/ves*. This parameter is subject to glucocorticoids regulation [31].
- *Number of cells N*_*a*_: we have not found any information about corticotroph number in human pituitary, only percentages of total pituitary cells are known approximately (*N*_*a*_/*N* ≈ 0.15 ~ 0.2) [117]. In the young rat (30 days) pituitary *N*_*a*_ ≈ 9×10^4^ cells are observed to secrete ACTH (figure 1 in [158]) of a total of approximately 1.4×10^6^ pituitary cells [104], which is consistent with reported percentages (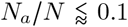 [158]). Therefore, we can only conclude that in human the number of corticotrophs must be higher than in rat (*N*_*a*_ > 1×10^5^). In addition, this number is modulated by both CRH stimulation and glucocorticoid regulation [31, 158, 104].

**Readily-Releasable Vesicle Pool replenishement:** using evidence according to which the time-constant for replenishement of the readily releasable vesicle pool is of *τ* ≈ 42 *sec* (rat), ≈ 10|20 sec (mouse, bovine) (observed *in vitro*) [165], we can estimate *K*_*smax*_ ≈ 1/*τ* in the equation describing the replenishement process ((51)). That is, assuming no secretion 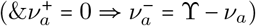, that all available vesicles have been exhausted (*ν*_*a*_(0) = 0); approximating the first Hill function term in equation (51) a with step function 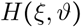 (where 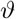 is a threshold, function of *θ* and *K*_*mv*_), and assuming that at basal activity 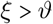 (so that vesicle synthesis is activated) we have:

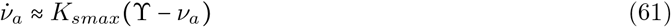

Therefore, *K*_*smax*_ ≈ 1/40 to 1/10 (*sec*)^−1^ (i.e. 1.42 to 6 (*min*)^−1^), essentially instantaneous at 1 *min*

**Table 3:**
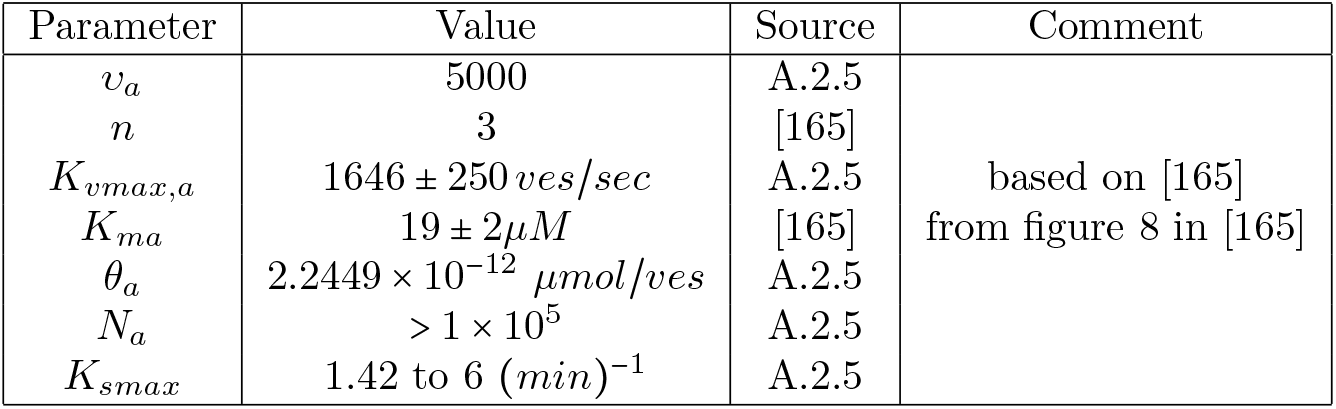
Estimated parameter typical values for the corticotroph secretion functional unit, *ν*_*a*_: vesicle number/cell, *n*: number of *Ca*^++^ ions binded, *K*_*vmax,a*_: maximum exocytosis rate, *K*_*ma*_: half maximal rate of exocytosis, *θ*_*a*_: ACTH availability/vesicle, *N*_*a*_: number of corticotrophs

## A.3 Ion channel models and electrophysiology

## A.3.1 Cell membrane

**Ion Channels:**

- Binding constants: *BKchannelCa*^++^ high affinity [157] in corticotrophs [148].
- *V*_50_ for BK STREX-1 variant channel exposed to 0.1 *μM*[*Ca*^2+^]_*i*_ in physiological potassium gradients: 30.7 ± 1.6 *mV* [148] 26.3 ± 5.8 *mV* in mouse AtT20 BK channels

**Corticotroph membrane potential:** following [97, 98, 150], we may define the change in membrane potential as:

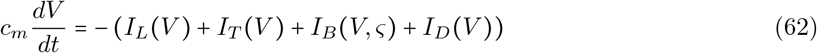

where *L* stands for L-type VGCC, *T* for T-type VGCC, *B* for Big-K channel, *D* for Delay-Rectifier-K; *c*_*m*_ is the membrane capacitance, *V* the membrane potential, *ς* = [*Ca*^++^]_*i*_ and *I*_*x*_ the ionic currents, for *x* being the L and T type VGCC currents, the Big conductance voltage and calcium sensitive currents and the delay rectifyer *K*^+^ currents. The currents are given by:

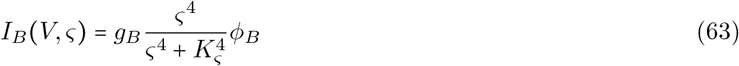

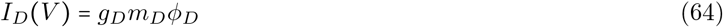

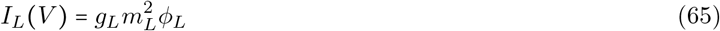

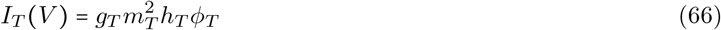

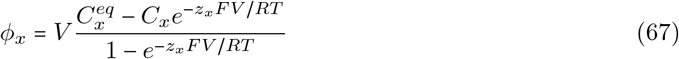

where *m*_*x*_ are activation variables whose steady states are given by 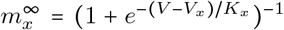 for *x* ϵ {*B, D, L, T*}, *C*_*x*_ and 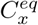 are, respectively, the transitory and equilibrium ion concentrations for each channel, and ς = [*Ca*^++^]_*i*_.

Several simplifications and modifications can be made to the formulation given in [97, 98], based on the following facts:

1. Channels activation/inactivation occurs instantaneously for the time-scale of interest (msec compared to sec,min).
2. Steady state activation/inactivation follows a steep squared Hill function with respect to V [97].
3. T-VGCC current magnitudes are negligible compared to other currents at steady-states.
4. L-VGCC current is almost insensitive to [*Ca*^++^]_*i*_ changes (physiological values of [*Ca*^++^]_*i*_ << [*Ca*^++^]_*e*_) and [*Ca*^++^]_*e*_ are assumed to be constant (see figure 12).
5. L-VGCC current dependence on membrane potential is almost symmetric with respect to *V* = 0 (for the parameters given in [97, 98], (see figure 12).
6. L-VGCC currents is always negative for physiological values of the variables involved (see figure 12).
7. DR and BK maximum magnitudes are very similar
8. DR currents show a first strong inflexion point (at ≈ −30*mV*) before which their almost zero and after which their are almost linear with respect to V and very steep, until a second not so strong inflexion point at (at ≈ −10*mV*) after which it is again almost linear with a slower slope (see figure 12).
9. BK currents follow an almost linear function with respect to V, with a similar slope to that shown by DR currents when the membrane polarization is low.

**Figure 12:**
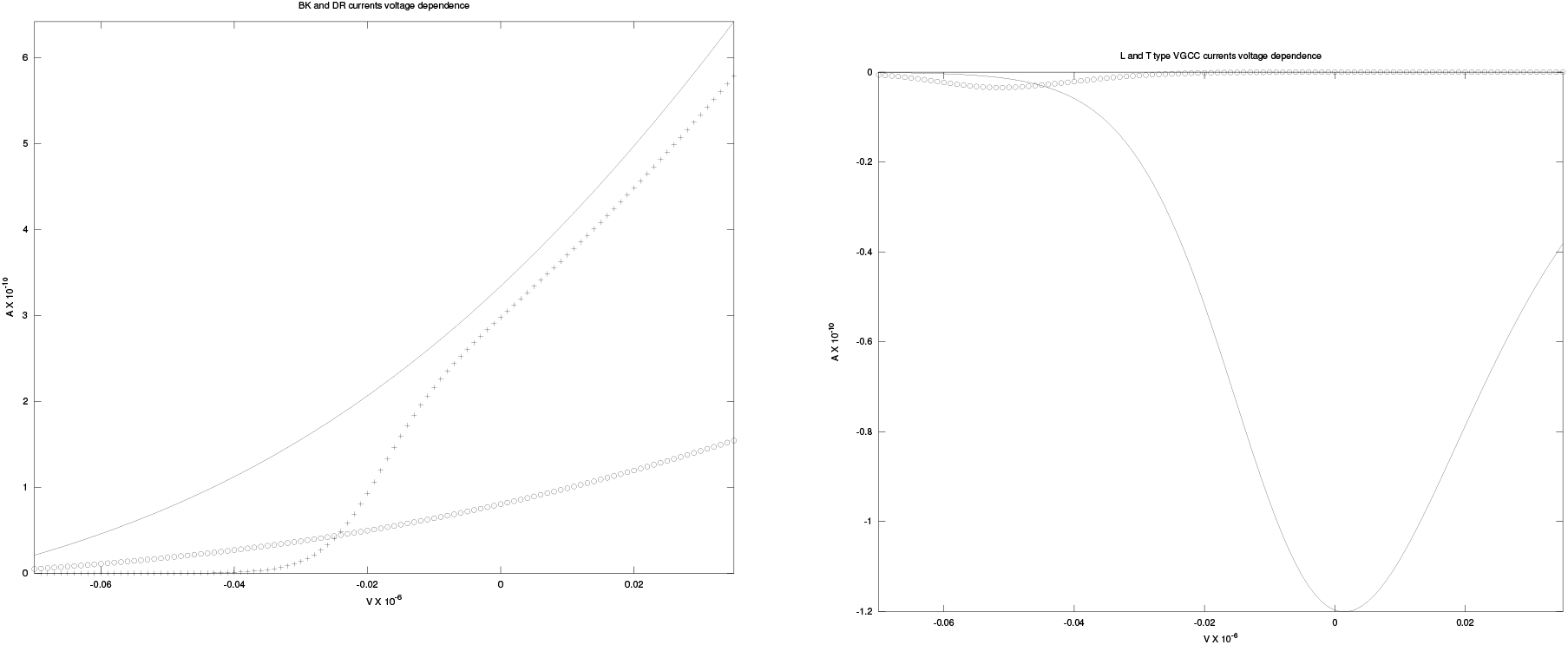
Voltage dependence of (left:) K-DR (+) and BK currents (for [*Ca*^++^]_*i*_ = 2 *μM* (line) and 0.3 *μM* (0)) and (right:) L (line) and T (o) type *Ca*^++^ currents, computed with steady-state current expressions from equations in [97, 98] and parameters given therein.

**Figure 13:**
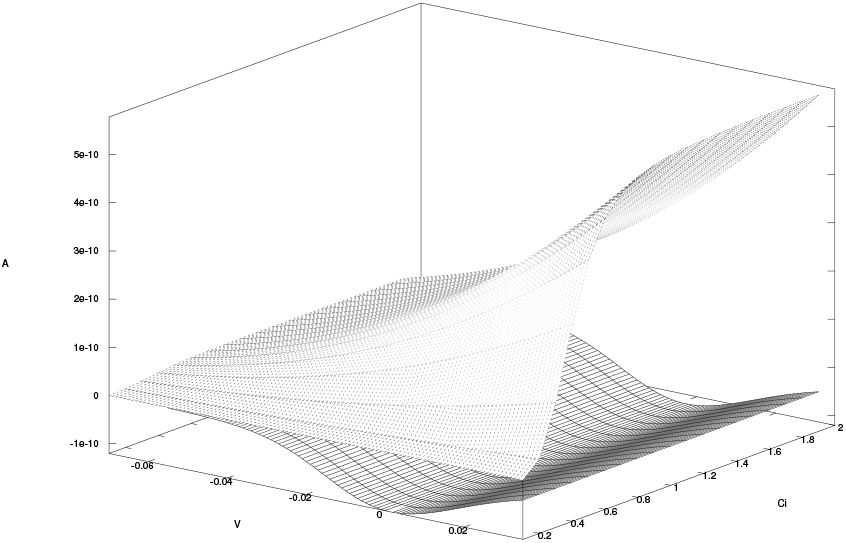
Superposition of BK and L-VGCC currents as a function of V and [*Ca*^++^]_*i*_, computed with steady-state current values from equations in [97, 98] and parameters given therein.

**Figure 14:**
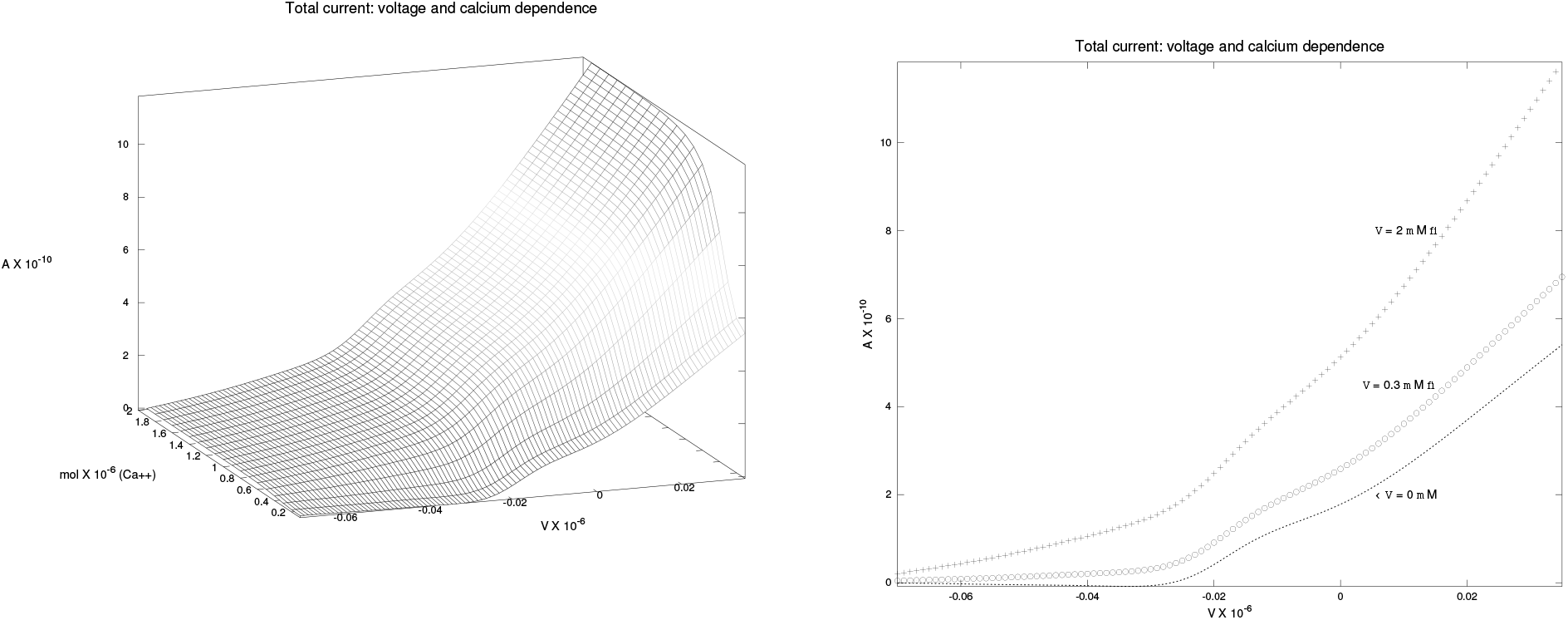
Total steady state ionic current sum across the membrane for different values of [*Ca*^++^]_*i*_, computed from asymptotic formulation of equations in [97, 98] and parameters given therein (Right: ‘+’: 2 *μM*, ‘o’: 2 *μM* 0.3 *μM*, ‘−’: 0 *μM* (null BK current)).

**Figure 15:**
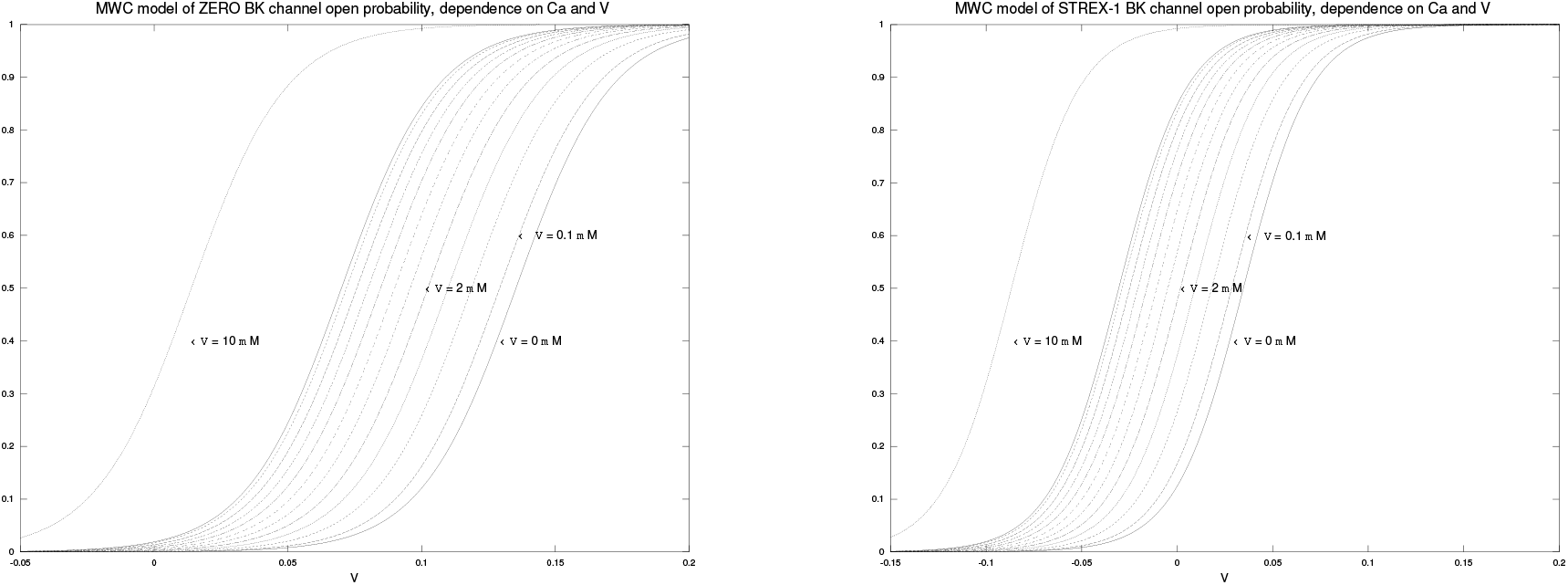
*BK channels model*. Left: open probability of BK channels as a function of membrane potential for different values of [*Ca*^++^]_*i*_, according to allosteric MWC model of *mslo* BK channel clone in [37]; this curves fit well with *V*_0.5(*max*)_ values for [*Ca*^++^]_*i*_ ∈ 0.1, 1, 10 *μM* and with the normalized conductance plot at [*Ca*^++^]_*i*_, = 1 *μM* for the ZERO type BK channel [26]; the fit is better for very low values of [*Ca*^++^]_*i*_] as explained in [37]. Right: Same model modified to account for STREX-1 variant lower voltage triggering, for *L* (0) = 7 fits *V*_0.5_ ≈ 30 *mV* for [*Ca*^++^]_*i*_ = 0.1 *μM* as given in [148].

Because of 1, we may use the steady-state form of the activation variables, neglecting very rapid dynamics.

In addition, an alternative and more precise formulation for BK channel activity was proposed by [37] that faithfully fits experimental data, the function is still sigmoid and steep and its parameters have been estimated in [37] for ZERO type BK channels; STREX-1 type channels that are the ones expressed in corticotrophs have a lower voltage gating characteristics [148], the shift of the open probability curve to lower voltages is mainly obtained by varying a single parameter (*L*(0): the the open-toclosed equilibrium constant in the absence of bound Ca^++^ at 0 *mV* [37]). Using the voltage values for half activation (*V*_50_) given in [148], an estimate of *L*(0) can be found by simply evaluating the activation function, a value of *L*(0) ≈ 7 has been found for the value *V*_50_ ≈ 30 *mV* given in [148].

The membrane potential at the time scale of interest (seconds-minutes) is therefore given by:

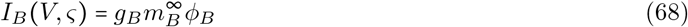

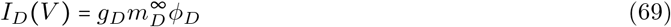

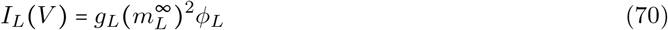

for 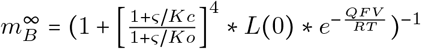

**Table 4:** Educated guesses of parameter values for STREX-1 variant of BK channel.

| Parameter | Value | Source | Comment |
| --- | --- | --- | --- |
| $L(0)$ | 7 | this work | Estimated such that $V_{50}(0.1 nM) \approx 30 mV$ for STREX-1 variant ([148]) |
| $K_o$ | 1 $\mu M$ | [37] | estimation for ZERO variant BK channel |
| $K_c$ | 10 $\mu M$ | [37] | estimation for ZERO variant BK channel |
| $Q$ | 1.4 | [37] | estimation for ZERO variant BK channel |

## Footnotes

1 [*Ca*^++^]_*e*_ = extracellular calcium

2 [*Ca*^++^]_*e*_ = intracelluar calcium stores

3 and that other cations, *Mg*^++^ and *Zn*^++^ are also involved in BK channel activation

5 Given that cAMP synthesis and degradation is very fast (seconds) [4], we may also assume that the steady-state is reached faster than the time resolution of interest, and we may replace the first reaction by a static transfer function corresponding to this experimental steady-state acummulation function

